# U2AF regulates the translation and localization of nuclear-encoded mitochondrial mRNAs

**DOI:** 10.1101/2024.09.18.613780

**Authors:** Gloria R. Garcia, Murali Palangat, Josquin Moraly, Benjamin T. Donovan, Bixuan Wang, Ira Phadke, David Sturgill, Jiji Chen, Brittney Short, Gustavo A. Rivero, David M. Swoboda, Naomi Taylor, Kathy McGraw, Daniel R. Larson

## Abstract

The mechanisms underlying molecular targeting to mitochondria remain enigmatic, yet this process is crucial for normal cellular function. The RNA binding proteins U2AF1/2 form a heterodimer (U2AF) that shuttles between the nucleus and cytoplasm, regulating splicing in the nucleus and translation in the cytoplasm. Our study identifies an unexpected role for U2AF in mitochondrial function. We demonstrate that U2AF interacts with nuclear-transcribed mitochondrial mRNAs and proteins, inhibits translation, localizes to the outer mitochondrial membrane, and regulates mRNA localization to mitochondria. Moreover, an oncogenic point-mutation in U2AF1(S34F) disrupts this regulation, leading to altered mitochondrial structure, increased translation, and OXPHOS-dependent metabolic rewiring, recapitulating changes observed in bone marrow progenitors from patients with myelodysplastic syndromes. These findings reveal a non-canonical role for U2AF, where it modulates multiple aspects of mitochondrial function by regulating the translation and mitochondrial targeting of nuclear-encoded mRNAs.

## Introduction

U2AF1 and U2AF2 are members of the serine- and arginine-rich (SR) family of proteins. SR proteins are RNA-binding proteins that regulate various aspects of mRNA metabolism, ranging from splicing to decay. Notably, a subset of SR proteins shuttle between the nucleus and cytoplasm, coordinating post-transcriptional steps of mRNA metabolism.^1^ U2AF1 and U2AF2 form a heterodimer (U2AF) that also shuttles and is bound to RNA in both cellular compartments. In the nucleus, U2AF regulates spliceosome assembly and 3’ splice site selection.^2^ Splicing-independent roles for U2AF have also been observed in RNA export and translational regulation^3–5^, similar to other SR proteins.^6,7^ However, the fact that RNA binding proteins (RBPs) have multiple overlapping roles in gene regulation -- often mediated by the same binding domain -- remains an obstacle in dissecting such regulation.^8^ Consequently, the physiological roles of SR proteins in coordinating post-transcriptional regulation remains elusive.

One approach to addressing this problem is through human genetics and the characterization of functional mutations in RNA-binding domains. For example, germline mutations in RBPs such as DBR1, TDP-43, FMRP, EWSR1, and DDX41 have provided unanticipated insights into the diverse roles of these RBPs.^9–11^ Similarly, a recurrent somatic point mutation in the first Zn-finger domain of U2AF1(S34F) has been identified across a range of cancers, including myelodysplastic syndrome (MDS), acute myeloid leukemia (AML), uterine carcinosarcoma^12^, and lung adenocarcinoma.^13^ This U2AF1-S34F mutation disrupts normal mRNA binding specificity, alters splicing kinetics, modulates R-loop abundance, and affects translational regulation both specifically and globally.^4,14–18^ The mutation was first reported in MDS, a group of clonal stem cell malignancies characterized by bone marrow failure, resulting in dysplasia, peripheral cytopenias, and a high risk of progression to AML.^19,20^ In MDS, the U2AF1-S34F mutation emerges early in disease development and is believed to contribute to the ineffective hematopoiesis characteristic of the disorder.^21–23^ Notably, mitochondrial dysfunction is a defining phenotype of MDS^19^, and splicing factor mutations are a defining genotype: >50% of MDS patients harbor mutations in U2AF1, SRSF2, ZRSR2, or SF3B1 within their hematopoietic stem and progenitor cells (HSPCs).^20,24^ Overall, the molecular mechanisms by which mutated U2AF1 drives cancer progression remain poorly understood, suggesting a possible splicing-independent function.

Here, we identify an unexpected direct role for U2AF in regulating mitochondrial function. Over 95% of mitochondrial proteins are encoded by nuclear genes and translated by cytosolic ribosomes, making molecular targeting critical for mitochondrial function.^25^ Targeting is primarily achieved via the peptides N-terminal mitochondrial targeting sequence (MTS), whose charge, structure, and size all contribute to protein import.^26^ The canonical mechanism involves mitochondrial proteins translated in the cytosol being maintained in an unfolded state by chaperones before being targeted to the mitochondrial translocase.^27^ Another mechanism, co-translational import via ribosomes docked on the mitochondrial surface, has also been reported.^26^ In addition to these mechanisms, RBPs have also been implicated in mitochondrial targeting.^28^ For instance, in budding yeast, RBPs bind specific sites in the 3’ untranslated region (UTR) of mRNAs to facilitate targeting to mitochondria. Recent transcriptome-wide studies in yeast and human cells suggest that translation and RNA localization work in concert in an mRNA-specific manner.^29,30^ However, the extent to which RBPs are involved in achieving this balance remains largely unknown. Moreover, it remains unclear how the targeting of molecules—such as mRNAs, ribosome-nascent chain complexes, and precursor proteins—to the mitochondria are altered during oncogenesis.

Using a combination of biochemical, sequencing, and advanced microscopy techniques in isogenic non-transformed cell lines and primary human bone marrow samples from MDS patients, we provide evidence that U2AF interacts with hundreds of nuclear-encoded mitochondrial mRNAs and proteins and is localized to the mitochondrial membrane. Using ribosome footprinting and in vitro reconstitution, we show that U2AF directly impedes translation in the N-terminal mitochondrial targeting sequence (MTS) region and concurrently regulates the targeting of mRNAs to mitochondria in single cells—a regulation crucial for maintaining proper mitochondrial structure and function. Notably, the U2AF1-S34F mutation significantly disrupts this regulation, leading to metabolic rewiring in hematopoietic stem and progenitor cells. These findings reveal an important role for the RNA binding protein U2AF in regulating the cross-talk between nuclear and mitochondrial genomes.

## Results

### U2AF interacts with mitochondrial mRNAs and proteins

To identify cytoplasmic proteins interacting with U2AF, we tagged U2AF1 with a 3XFLAG epitope (**Fig. 1A**) and validated genomic integration and binding to U2AF2 in human bronchial epithelial cells (**Fig. 1B and Fig. S1A**). We performed subcellular fractionation on parental and *U2AF1^wt/wt-3XFLAG^* cell lines to obtain cytoplasmic and nuclear/ER fractions (**Fig. S1B-C**), followed by 3XFLAG immunoprecipitation and TMT-labelled quantitative mass spectrometry. The parental cell line served as a non-specific background control. Analysis of the cytoplasmic fraction revealed U2AF1 and U2AF2 as top hits, indicating high data quality, and highlighted significant enrichment of U2AF1-interacting proteins associated with mitochondrial functions (**Fig. 1C and Table S1**). Gene set enrichment analysis (GSEA) showed enrichment for ‘Splicing,’ aligning with the role of U2AF in the nucleus, and ‘Oxidative Phosphorylation’, revealing an unexpected cytoplasmic function (**Fig. 1D and Fig. S1D**).

**Fig. 1.**
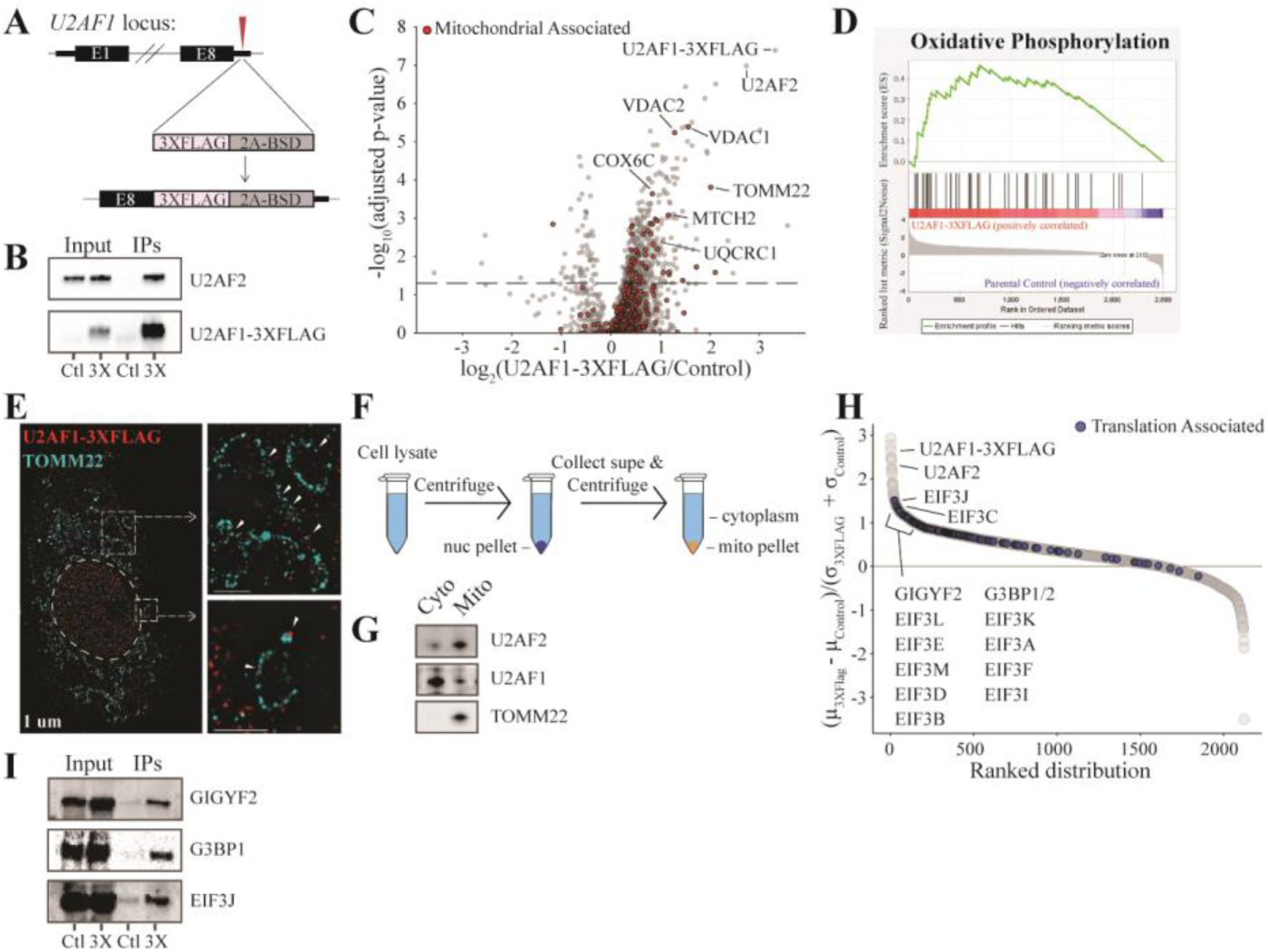
Cytoplasmic U2AF1 interacts with mitochondrial proteins and localizes at the outer mitochondrial membrane. (**A**) Schematic for CRISPR-Cas9-mediated insertion of a C-terminal 3XFLAG tag into the *U2AF1* gene locus. (**B**) Validation of U2AF1 endogenous 3XFLAG tagging by immunoprecipitation and western blot in whole cell lysates from parental (Ctl) and U2AF1 edited (3X) cell lines. (**C**) Volcano plot analysis of the cytoplasmic fraction from parental and *U2AF1^wt/wt-3XFlag^* cell lines. Mitochondrial-associated proteins are highlighted in red. A dashed line denotes the threshold for statistical significance (adjusted p < 0.05). Data represent three biological replicates (n=3) for each sample group. (**D**) GSEA plot showing significant enrichment of the ‘Oxidative Phosphorylation’ category in *U2AF1^wt/wt-3XFlag^* cells compared to the parental control, as determined by the KEGG gene set. (**E**) Super-Resolution TIRF microscopy visualization using DNA-PAINT in *U2AF1^wt/wt-3XFlag^* cells. U2AF1 was labeled with an anti-FLAG antibody (red), and the outer mitochondrial membrane was identified with anti-TOMM22 (cyan). Insets show magnified areas highlighting the colocalization of U2AF1 with the mitochondrial outer membrane. Scale bar indicates 1 µm. (**F**) Schematic of the centrifugation method used to separate mitochondrial (mito)-enriched and cytoplasmic fractions from cell lysates. (**G**) Western blot analysis of U2AF1, U2AF2 and TOMM22 (mitochondrial marker) in cytoplasmic and mitochondrial fractions from *U2AF1^wt/wt^ ^cells^*. (**H**) GSEA gene score plot of the nuclear-ER fraction from control and *U2AF1^wt/wt-3XFlag^* cell lines. Translation-associated proteins are highlighted in blue. Data represent three biological replicates (*n* = 3) for each sample group. (**I**) Western blot validation of U2AF1-interacting proteins GIGYF2, G3BP1, EIF3(J subunit) from the ER-nuc fraction of parental (Ctl) and U2AF1 edited (3X) cell lines.

To assess U2AF localization at mitochondria, we used imaging and cellular fractionation. First, we implemented super-resolution TIRF microscopy and DNA-PAINT labeling to co-stain U2AF1-3XFLAG and TOMM22, a mitochondrial outer membrane protein identified in our mass spectrometry data. Qualitative analysis showed U2AF1-3XFLAG predominantly in the nucleus with a diffuse cytoplasmic signal (**Fig. 1E**). Co-localization with TOMM22 confirmed its association with intact mitochondria, consistent with our mass spectrometry data, though U2AF is not exclusively mitochondrial. Second, we separated mitochondrial-enriched and cytoplasmic fractions from *U2AF1^wt/w^* cells using centrifugation and detected enrichment of U2AF in the mitochondrial fraction by Western blot analysis (**Fig. 1F-G**). Overall, three orthogonal approaches identified an interaction between the U2AF heterodimer and mitochondrial proteins.

In the nuclear-ER fraction, we identified an association between U2AF1 and proteins involved in translation (**Fig. 1H and Fig. S1E**). Notably, U2AF1 pulled down 12 of 13 subunits of the EIF3 translation initiation complex and ribosome quality control factors GIGYF2 and DDX6. Previous studies have implicated EIF3 in mitochondrial homeostasis.^31^ Several U2AF1-interacting proteins were validated by western blot (**Fig. 1I**). Our results suggest U2AF is involved in cross-talk between the nucleus and mitochondria, potentially at the level of translation and/or transport, hinting at a broader role in mitochondrial functions.

These protein-protein interactions prompted a re-assessment of U2AF1 protein-RNA interactions. From our previously published U2AF1 PAR-CLIP dataset^4^, we generated a stringent list of 2,736 U2AF1 cytoplasmic targets **(Table S2**). Meta-transcript analysis of all target mRNAs revealed a 5’ bias in U2AF binding, with peak enrichment just downstream of the translation start site, a region critical for targeting nascent proteins via the MTS (**Fig. 2A**). Next, we performed a Fisher’s exact test for association with mitochondrial mRNAs (**Fig. 2B)**. Results showed a significant 1.5-fold enrichment in U2AF binding to nuclear-encoded mitochondrial mRNAs (p < 0.0001) across key mitochondrial pathways (**Fig. 2C**). Examples include mitochondrial outer membrane translocase *TOMM70* and chaperone *HSPD1* (**Fig. 2D-E**). Next, we re-analyzed existing PAR-CLIP datasets for EIF3A, B, D, and G (combined as a single EIF3 track), G3BP2, and FMR1 using our computational pipeline (see methods).^32–34^ EIF3 and G3BP2 were identified in our nuclear/ER U2AF protein interactome, but FMR1 was not. Interestingly, Fisher’s exact test on the PAR-CLIP datasets showed a significant association of both EIF3 and U2AF with mitochondrial mRNAs, including co-association with the same mRNAs (**Table S3**). As control, FMR1 targets were not enriched with mitochondrial mRNA by this analysis. Thus, both protein-protein and protein-RNA interactions suggest a coordinated role between EIF3 and U2AF in regulating mitochondrial gene expression.

**Fig. 2.**
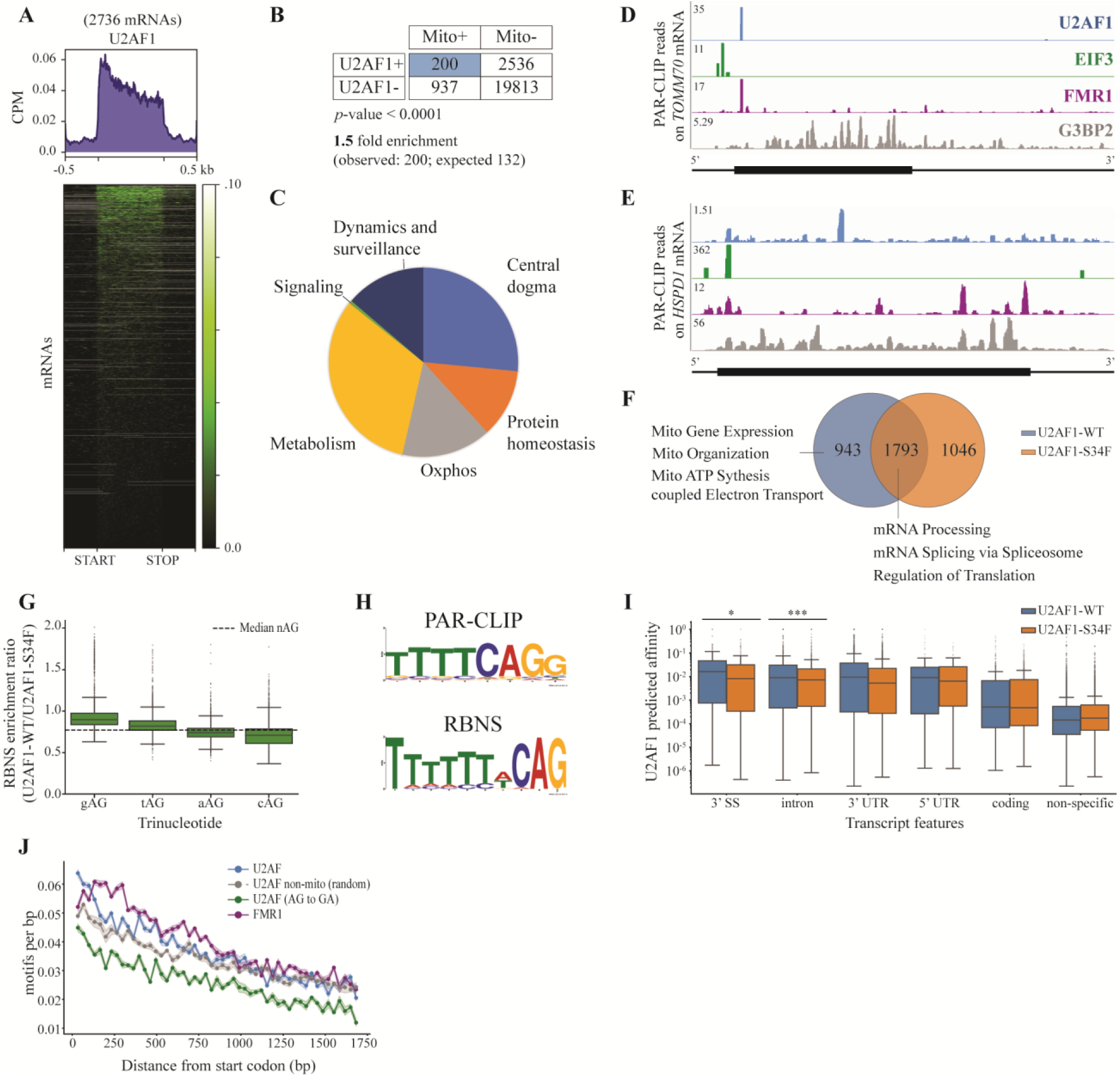
U2AF1 is bound to nuclear encoded mitochondrial mRNAs in the cytoplasm. (**A**) Meta-transcript analysis of U2AF1 binding on mRNAs, displaying normalized U2AF1 footprint reads across 2735 identified U2AF1 targets. Binding distribution is shown from 500 bp upstream to 500 bp downstream of the translation start site (START) to the stop codon (STOP). Gray lines in heatmap indicate absence of data. Nucleotide distances in kilobases (kb). (**B**) Fisher’s exact test results displayed in a contingency table, assessing the binding of U2AF1 to mitochondrial mRNAs, as defined by the MitoCarta 3.0 annotations. (**C**) Pie chart categorization of U2AF1-positive mitochondria-associated mRNAs using MitoCarta 3.0 pathway annotations. (**D-E**) Transcriptome browser tracks displaying PAR-CLIP binding profiles: cytoplasmic U2AF1 (blue), EIF3 (green), FMR1 (purple), and G3BP2 (gray) from whole cell extracts. (**D**) PAR-CLIP reads on *TOMM70* mRNA, with maximum read coverage shown on the right of the y-axis. (**E**) PAR-CLIP reads on *HSPD1* mRNA, similarly annotated for read coverage. (**F**) Venn Diagram of mRNA targets from PAR-CLIP datasets comparing U2AF1-WT (blue) and U2AF1-S34F (orange). The numbers represent the count of unique and shared mRNA targets. Significant functional enrichment terms for select biological processes are indicated for each group. (**G**) Comparative analysis of U2AF1-WT and U2AF1-S34F RNBS enrichment values for oligos containing a ‘nAG’ trinucleotide sequence. Median value for all trinucleotide sequences is indicated by horizontal dashed line on plot. Error bars denote the interquartile range. Outliers are represented as individual points. (**H**) Motif analysis comparing U2AF1-WT and -S34F binding preferences in PAR-CLIP (*in vivo*) and RBNS (*in vitro*) datasets. Using U2AF1-WT as the background model, enrichment analysis was conducted on U2AF1-S34F data to identify motifs preferentially enriched in the presence of the S34F mutation. (**I**) Boxplot of U2AF1-WT and U2AF1-S34F RBNS predicted affinity values assigned to respective *in vivo* PAR-CLIP binding sites. Transcript features—3’ splice site (3’ SS), intron, 3’ untranslated region (3’ UTR), 5’ UTR, coding regions, and non-specific sites—are displayed in descending order of their median RBNS-predicted affinity values for U2AF1-WT. Statistically significant differences between U2AF1-WT and U2AF1-S34F affinities are indicated (\**p* < 0.05 and \*\*\**p* < 0.001, Mann-Whitney U test with Benjamini-Hochberg correction). Error bars show the interquartile range, and points outside these bars are outliers. (**J**) 3’ Splice Site Motif Analysis in Mitochondrial and Non-Mitochondrial mRNAs. Frequency of 3’ splice site (3’ SS) motifs per base pair (bp) along the coding sequence relative to the start codon. Blue dots represent U2AF binding sites on mitochondrial mRNAs, while gray dots depict U2AF sites on non-mitochondrial (random) mRNAs of equal sample size (∼1,000 mRNAs), serving as a control. The green dots indicate U2AF binding sites on mitochondrial mRNAs where the AG dinucleotide in the 3’ SS motif has been swapped for GA. Purple dots represent FMR1 binding sites on mitochondrial mRNAs. The bin size for the analysis is 33 bp. The motif search was conducted using FIMO with a relaxed stringency (*p* < 0.01). Error bands indicate bootstrap error.

Understanding the mechanistic implications of RBP interactions remains a challenge in the field, especially considering the multiple overlapping roles that RBPs play in RNA homeostasis. As a perturbation to disrupt normal U2AF1 functions, we utilize the U2AF1-S34F mutation, identified as a heterozygous pathogenic mutation in blood diseases like MDS which disrupts U2AF1-mRNA interactions.^17,20^ We therefore analyzed cytoplasmic PAR-CLIP datasets to compare the mRNA binding profile of wild-type U2AF1 with its oncogenic S34F variant (**Table S4**). Meta-transcript analyses indicated similar binding profiles for U2AF1-WT and U2AF1-S34F across transcript regions, but with slightly altered positioning of U2AF1-S34F on transcripts (**Fig. S1A-B**). However, PAR-CLIP analysis of hits revealed a 47% overlap in mRNA targets between U2AF1-WT and the S34F mutant (**Fig. 2F**). Functional enrichment of the shared targets highlight their involvement in RNA processing, splicing, and translation. Yet, mRNA targets exclusive to U2AF1-WT showed a pronounced enrichment in mitochondrial processes, with these functional terms absent from the S34F dataset. This observation points to a specific role for direct interactions between U2AF1 and RNA binding sites in mitochondrial biology that is disrupted by the S34F mutation.

Next, we explored the nucleotide binding preferences of U2AF by combining PAR-CLIP (*in vivo* interactions) with RNA Bind-n-Seq (RBNS, *in vitro* affinity). RBNS relies on interaction between purified U2AF heterodimer from insect cells and a random library of 12-mer RNA oligos in solution. RNAs bound to U2AF are isolated by U2AF1-FLAG immunoprecipitation, and the resulting pull down is sequenced along with the input and free oligo pool. The affinity is determined with the Probound model, which takes as experimental input the relative fraction of a particular 12-mer oligo in each pool.^35^ Comparing the binding preferences of U2AF1-WT and S34F, we observed distinct sequence affinities at the typical U2AF1 binding site (-AG): ‘gAG’ sites showed the largest decrease in binding in S34F, while ‘cAG’ sites had increased binding (**Fig. 2G**). Interestingly, ‘gAG’ sites occur infrequently (< 1%) at annotated splice sites, indicating the largest changes in binding occur at sites which are virtually never used for splicing (**Fig. S1C**). U2AF1-S34F motif analysis using U2AF1-WT as the background model revealed ‘cAG’ enrichment in S34F, consistent with previous studies (**Fig. 2H).**^36^ Finally, we combined the PAR-CLIP and RBNS to obtain the predicted affinity of all *in vivo* sites for both U2AF1-WT and S34F. The strongest binding sites are indeed at the 3’ SS, with sites in the CDS almost an order of magnitude weaker but still well above the non-specific level (**Fig. 2I**).

Based on this finding of weaker sites in the CDS, we executed a straightforward motif search for the U2AF binding site in spliced mitochondrial mRNA at a relaxed stringency. Our motivation was to use an orthogonal analysis--independent of our experimental PAR-CLIP analysis– to identify the basis of U2AF specificity for these transcripts. For the motif, we used the canonical 3’ SS from the literature^37^ and searched in the coding sequences of all 1,136 mitochondrial genes from MitoCarta 3.0 using FIMO (**Fig. 2J**, FIMO, p<0.01). Indeed, we found a significant enrichment near the 5’ end (< 150 nt) of mitochondrial mRNA compared to control analysis in mRNA coding for random groups of transcripts, recapitulating our empirical binding data (**Fig. 2A**). Swapping the AG dinucleotide for GA in the motif search reduced the difference between mitochondrial transcripts and random control, indicating the canonical U2AF binding sequence is important and not just the sequence content. Additionally, the same analysis for FMR1 showed a difference in binding but not in the MTS region. Thus, the specificity for U2AF interaction with mitochondrial transcripts is encoded in the genome.

In summary, U2AF is bound to spliced transcripts in the cytosol which are enriched for mRNAs encoding mitochondrial proteins. These binding sites are preferentially located in the CDS downstream of the translation start site and are weaker compared to those at 3’SS. Nevertheless, we are able to enrich these sequences in RBNS, measure changes in affinity in the presence of the S34F mutation, and identify a sequence feature computationally.

### U2AF binding regulates translation and localization of nuclear-encoded mitochondrial RNA

The observation that U2AF binds to a region of the CDS that is critical for early elongation and protein targeting^31,38^ led us to investigate how U2AF might directly influence translation. First, we aimed to map ribosome positions relative to U2AF binding sites. Using polysome profiling followed by Western blot analysis on the *U2AF^wt/wt-3XFlag^* cell line (**Fig. S3A**), we found that U2AF1-3XFLAG is mainly associated with mRNA with few or no ribosomes. We then refined existing methods to measure single ribosomes (monosomes) and closely spaced ribosomes (disomes)^39^ with improved small RNA library preparation (see methods, **Fig. S3B-D**).^40^

To elucidate the sequence-dependent relationship between RBP binding sites and ribosome positions on mRNAs, we performed cross-correlation analysis between Ribo-seq and PAR-CLIP coverage plots for several RBPs (U2AF1, EIF3, G3BP2, FMR1) on 2,736 U2AF targets in WT (*U2AF^wt/wt^*) or heterozygous mutant (*U2AF^w/tS34F^*) cells (**Fig. 3A-B**).^41^ Cross-correlation is computed as the normalized cross-covariance over all possible positions using an incrementally increasing binsize. We use only the T to C reads in the PAR-CLIP data, which are the reads most closely associated with direct interaction between the RPB and the sequence. Importantly, this analysis takes the sequence reads as direct input, without the need for peak calling or additional filtering. The displayed result is oriented relative to the RBP: the RBP is position zero, with positive correlation corresponding to ribosome footprints which are located in the 3’ direction, and negative correlation corresponding to ribosome footprints in the 5’ direction.

**Fig. 3.**
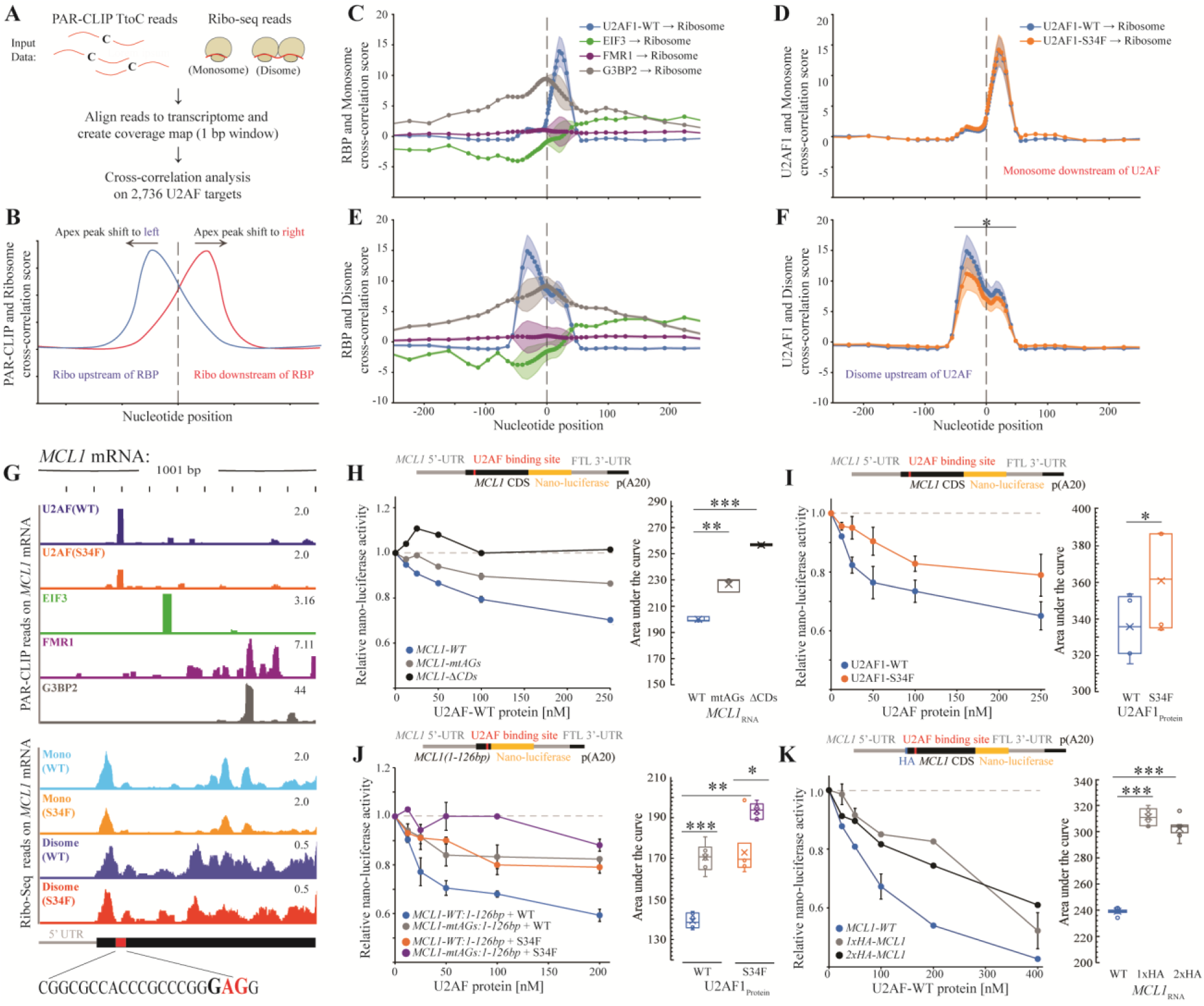
U2AF1 impedes translation in a sequence-, position-, and RNA binding protein-dependent manner. (**A**) Schematic of the cross-correlation analysis pipeline. (**B**) Diagram illustrating the interpretation of cross-correlation plots. (**C**, **E**) Cross-correlation analysis between RBP and monosome (**C**) or disome (**E**) ribosome footprints. U2AF1-WT (blue), EIF3 (green), FMR1 (purple), and G3BP2 (grey). Error bands represent the standard error of the mean (SEM). (**D**, **F**) Cross-correlation analysis between U2AF1 (WT and S34F mutant) and ribosome footprints (monosome and disome). U2AF1-WT (blue) and U2AF1-S34F mutant (orange). Error bands represent the standard error of the mean (SEM). Statistical significance was determined using a *t*-test. (**G**) Browser tracts showing PAR-CLIP reads for U2AF1 (WT and S34F), EIF3, FMR1, and G3BP2 on the *MCL1* mRNA. Ribo-seq reads for monosomes and disomes (*U2AF1^wt/wt^* and *U2AF1^wt/s34F^*) are also shown. The main U2AF binding site is highlighted, with a sequence detail of the binding motif (**GAG**). (**H**) *In vitro* translation reporter assay using a capped Nano-Luciferase reporter mRNA with the *MCL1* coding sequence. Dose-response plot shows relative nano-luciferase activity in the presence of increasing concentrations of U2AF-WT protein. Constructs tested include *MCL1-WT*, *MCL1-mtAGs* (mutated U2AF binding site), and *MCL1-ΔCDS* (coding sequence deletion). The right panel shows the area under the curve (AUC) analysis to highlight significant differences. For all *in vitro* assays shown, error bars represent the standard error of the mean (SEM) and statistical significance was determined using a t-test. \**p* < 0.05, ** *p* < 0.01, \*\*\**p* < 0.001. (**I**) Dose-response plot of the *MCL1 in vitro* translation reporter assay comparing U2AF1-WT and U2AF1-S34F mutant proteins. (**J**) *In vitro* translation reporter assay with *MCL1* constructs truncated downstream of the U2AF binding site. Dose-response plot shows relative nano-luciferase activity for *MCL1(1-126bp)* constructs with either WT or mutated AG binding sites (mutated U2AF binding site), in the presence of U2AF-WT or U2AF1-S34F proteins. (**K**) *In vitro* translation reporter assay testing the impact of a HA tag insertion immediately downstream of the start codon on *MCL1* translation. Dose-response plot shows relative nano-luciferase activity for *MCL1-WT*, *1xHA-MCL1*, and *2xHA-MCL1* constructs in the presence of the U2AF-WT protein.

This correlation analysis indicates that U2AF binding is correlated with upstream disome accumulation. In *U2AF^wt/wt^* and *U2AF^wt/S34F^*, monosomes were more likely positioned in a discrete region ∼30 nt downstream of U2AF1 sites (**Fig. 3C-D**). EIF3 sites correlate with a broad distribution of monosomes downstream, consistent with its role as a translation initiation factor.

G3BP2 exhibited a more symmetric distribution, and FMR1 showed minimal cross-correlation on these transcripts. In contrast, disomes had a higher probability of being located immediately upstream of U2AF1 sites, suggesting U2AF1 may influence ribosome pausing or stalling (**Fig. 3E**). Significant differences were observed between *U2AF^wt/wt^* and *U2AF^wt/S34F^* in the peak region (nucleotide positions −50 to +50), with a reduction in disome formation at U2AF1 binding sites in *U2AF^wt/S34F^* (**Fig. 3F**). EIF3 displayed disomes downstream similarly to monosomes; G3BP2 was largely symmetric, and FMR1 showed a flat correlation (**Fig. 3E**). Thus, U2AF1 appears to exhibit a unique positional relationship with ribosomes on these target transcripts compared to other RBPs. These data, combined with U2AF1 location in the polysome profile, suggest that bound U2AF1 acts as a barrier to translation, and in the presence of the S34F mutation, this barrier is reduced, resulting in less ribosome stalling and potentially more efficient elongation.

To directly test the model that U2AF1 affects the translational output of a nuclear-encoded mitochondrial gene, we implemented a U2AF-dependent *in vitro* translation assay. This assay included: 1) a capped Nano-Luciferase reporter mRNA fused to the *MCL1* coding sequence, 2) translation-competent HeLa cell cytosolic extracts, and 3) purified U2AF heterodimer with RNA-binding activity from insect cells (**Fig. S3E-F)**. We chose *MCL1* because it codes for a mitochondrial protein with a strong PAR-CLIP peak in the 5’ CDS containing a gAG motif which shows the largest changes in binding by our RBNS data. Moreover, both adenosines in the site can be mutated without altering the amino acid sequence (**Fig. 3G and Fig. S3E**). Finally, *MCL1* has direct disease relevance in myeloid malignancies as an anti-apoptotic member of the *BCL* family which confers resistance to BCL-2 inhibition by ventoclax.^42^

We found that U2AF1 functions as a translational repressor for *MCL1* mRNA in a sequence-, position-, and RBP-dependent manner. First, U2AF1-WT inhibited translation in a concentration-dependent manner (**Fig. 3H**). Mutation of the U2AF binding site (*MCL1-mtAGs*) significantly reduced this inhibition, and deletion of the *MCL1* coding sequence (*MCL1-ΔCDS*) completely abolished it, indicating that U2AF1 directly represses translation through interactions with the *MCL1* CDS. Second, comparison between U2AF1-WT and U2AF1-S34F showed that the S34F mutant significantly alleviates translational repression (**Fig. 3I**). Third, truncation of the *MCL1* construct downstream of the U2AF1 binding site, retaining only the first 126 bp (*MCL1(1-126bp)*), continued to exhibit translational repression by U2AF1-WT (**Fig. 3J**). Moreover, this minimal construct was sensitive to both *cis*-acting RNA mutations and *trans*-acting protein mutations. In contrast, FMR1, another RNA-binding protein known to alter translation elongation (albeit not on this transcript), did not significantly reduce translation at comparable concentrations (25 and 50 nM).^43^ However, it did inhibit translation at higher concentrations (100-400 nM), presumably due to non-specific RNA binding (**Fig. S3G-H**). Previous publications suggest that the position of ribosome pausing relative to the start codon correlates with its impact on protein output, with pauses closer to the start codon exerting a stronger repressive effect.^44^ Specifically, it has been proposed that the strength and 5′ polarity of the pause, along with the translational efficiency of the gene, collectively determine the degree to which protein output is affected. To investigate whether the position of the U2AF binding site similarly influences its translational repression, we inserted HA tags immediately downstream of the start codon but upstream of the U2AF binding site (*1xHA-MCL1* and *2xHA-MCL1*). These modifications reduced the U2AF1 inhibitory effect on translation, indicating that the proximity of the U2AF binding site to the start codon is crucial for its repressive function (**Fig. 3K**). These *in vitro* data, coupled with *in vivo* ribosome footprinting and PAR-CLIP data, indicate that U2AF directly modulates translation in a region of the transcript critical for mitochondrial targeting.

Previous studies have shown a close coupling between translation, RNA localization, and mitochondrial protein targeting.^29,30^ To test if U2AF-induced changes in translation affect mRNA targeting, we analyzed the spatial localization of nuclear-encoded mitochondrial mRNAs relative to mitochondria using two-color smFISH in both *U2AF1^wt/wt^* and *U2AF1^wt/S34F^* cells. We examined six nuclear-encoded mitochondrial mRNAs: *TOMM6* (non-U2AF target control), *MCL1*, *TOMM20*, *HSPD1*, *HSPA9*, and *GHITM*, identified as U2AF targets by PAR-CLIP. Their localization to mitochondria was assessed using *MT-ND5*, an RNA encoded and transcribed within mitochondria (**Fig. 4A-F**).

**Fig. 4.**
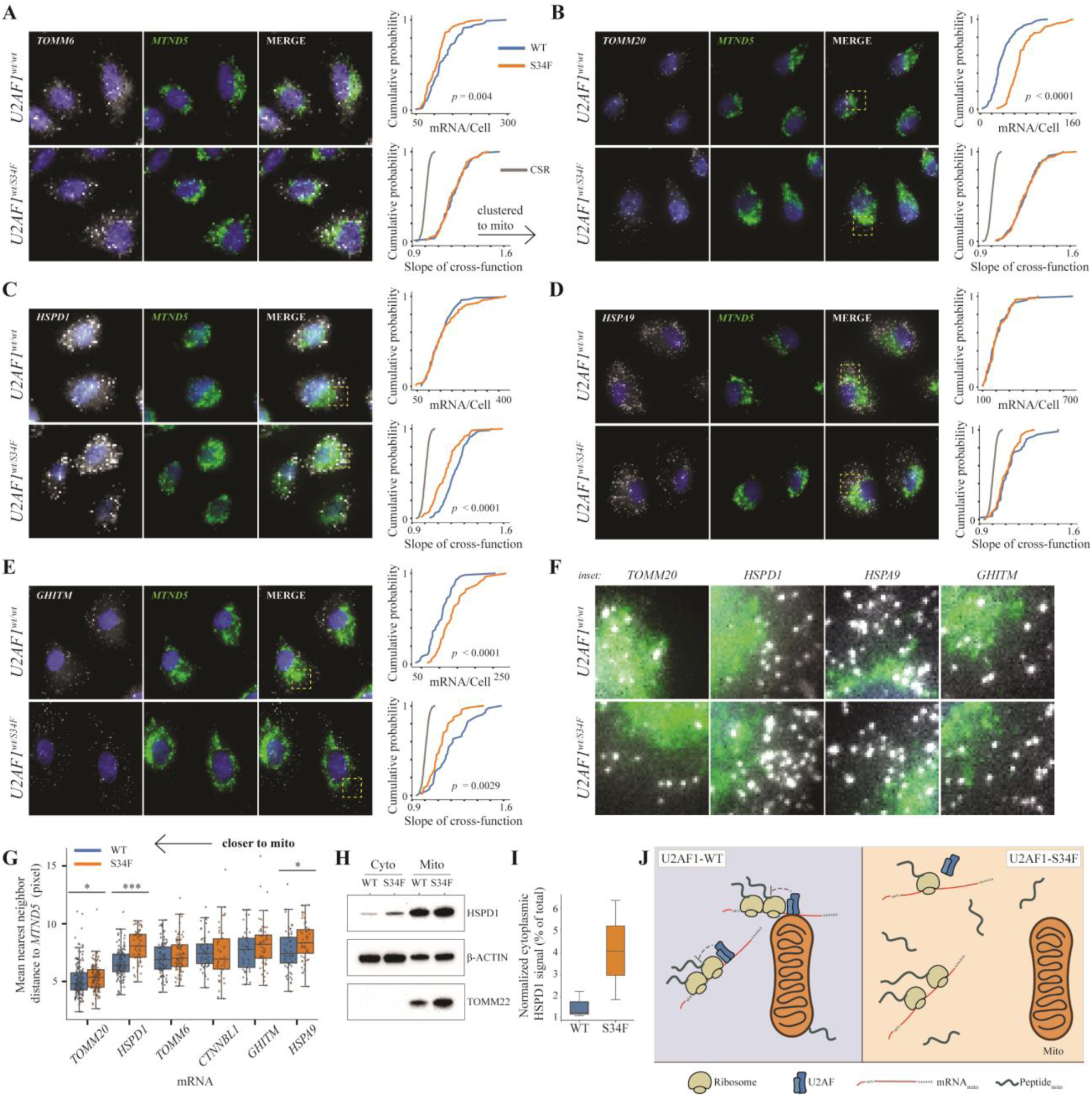
U2AF1 regulates mRNA localization and protein targeting to mitochondria. (**A-E**) Two-color smFISH analysis of nuclear-encoded mitochondrial mRNAs (*TOMM6*, *TOMM20*, *HSPD1*, *HSPA9*, *GHITM*) and the mitochondrial marker *MTND5* in *U2AF1^wt/wt^* and *U2AF1*^wt/S34F^ cells. The right-side plots show the cumulative distribution function (CDF) of mRNA per cell (top) and the CDF of the slope of the Cross-K function (bottom). The Cross-K function analyzes the spatial relationship between all mRNA and mitochondrial signals within the cell (global analysis), with a gray line indicating completely spatially random (CSR) simulated data. An arrow on the Cross-K function plots indicates that data shifted to the right represent mRNA patterns more clustered to the mitochondrial signal. Significance was tested with a Mann-Whitney U test, and p-values are shown if significant. *TOMM6* is a non-U2AF target control, while *TOMM20*, *HSPD1*, *HSPA9*, and *GHITM* are U2AF targets according to PAR-CLIP data. (**F**) Zoomed-in images (inset) of *TOMM20*, *HSPD1*, *HSPA9*, and *GHITM* mRNAs in *U2AF1^wt/wt^* and *U2AF1^wt/S34F^* cells, outlined with a dotted yellow box in panels A-E, showing the mRNA localization relative to *MTND5*. (**G**) Mean nearest neighbor distance analysis of nuclear-encoded mitochondrial mRNAs (*TOMM6*, *TOMM20*, *HSPD1*, *HSPA9*, *GHITM*) and non-mito control mRNA (*CTNNBL1*) relative to *MTND5* in *U2AF1*^wt/wt^ and *U2AF1^wt/S34F^* cells. Data are shown as rank-ordered box plots with significance tested using a Mann-Whitney U test. \**p* < 0.05, ** *p* < 0.01, \*\*\**p* < 0.001. (**H**) Western blot analysis of HSPD1, β-ACTIN (loading control), and TOMM22 (mitochondrial marker) in cytoplasmic and mitochondrial fractions from *U2AF1^wt/wt^* and *U2AF1^wt/S34F^* cells. (**I**) Quantification of HSPD1 protein levels in the cytoplasmic fraction from three independent experiments, normalized to β-ACTIN signal. Data is presented as box plots with significance tested using a Mann-Whitney U test. (**J**) Schematic representation illustrating the proposed mechanism via which U2AF1-WT and U2AF1-S34F impact mitochondrial mRNA translation/localization and peptide targeting to the mitochondria.

We employed two complementary spatial analysis methods: the Cross-K function, which evaluates mRNA clustering around mitochondria (*MT-ND*5), and the mean nearest neighbor distance, which measures mRNA proximity to mitochondria (see methods). Importantly, we note there are clearly both co-localized and non-colocalized transcripts, necessitating this statistical analysis. *TOMM6*, the non-U2AF target, showed no significant difference in spatial pattern or distance to the *MTND5* signal between *U2AF1^wt/wt^* and *U2AF1^wt/S34F^* cells, indicating that observed changes are specific to U2AF binding (**Fig. 4A**). The *MCL1* smFISH probes yielded a poor signal-to-noise ratio, making them unsuitable for spatial analysis due to inadequate spot detection. All other U2AF target mRNAs in the *U2AF1^wt/S34F^* line showed significant disruptions in mRNA targeting to the mitochondria, with mRNAs being less clustered around and/or further from the mitochondrial signal (**Fig. 4C,E,G**). These results are consistent with our model that U2AF1-mediated translational regulation facilitates efficient mRNA targeting to mitochondria, which is disrupted by the S34F mutation.

Subsequently, we examined the consequences at the protein level, specifically investigating whether HSPD1 protein targeting to mitochondria was disrupted in *U2AF1^wt/S34F^* cells. HSPD1 was chosen because smFISH showed no changes in mRNA abundance per cell between *U2AF1^wt/wt^*and *U2AF1^wt/S34F^* cells but significant differences in mRNA clustering and distance to mitochondria. Using a mitochondrial fractionation assay as described in (**Fig. 1F**), we found approximately 3-fold more HSPD1 protein in the cytoplasmic fraction of *U2AF1^wt/S34F^* cells compared to *U2AF1^wt/wt^*cells (**Fig. 4H-I**). We also observed global increase in mitochondrial protein levels not reflected at the mRNA level in *U2AF1^wt/S34F^* cells (**Fig. S4A-C**). In summary, our results suggest that U2AF1 impedes translation of mitochondrial mRNAs, and the S34F mutation alleviates this block, leading to altered mRNA localization, increased mitochondrial protein levels, and changes in protein targeting (**Fig. 4J**).

### U2AF1-S34F mutation recapitulates metabolic remodeling seen in HSPCs from MDS patients

These data suggest an interplay between translation, localization, and targeting that may affect mitochondrial function. To determine whether U2AF regulates mitochondrial metabolism, we assessed the impact of OXPHOS on protein synthesis, one of the most energy-consuming intracellular processes.^45,46^ We used puromycin incorporation, an aminoacyl-tRNA analog that covalently attaches to C-terminus of nascent polypeptides, as a measure of translation and evaluated the importance of mitochondrial metabolism on translation by inhibiting mitochondrial ATP synthesis with oligomycin (**Fig. 5A**).^45^ Strikingly, under control conditions, puromycin incorporation was significantly higher in *U2AF1^wt/S34F^*cells compared to *U2AF1^wt/wt^* (**Fig. 5B-C**), indicating higher basal translation output and energy consumption in the *U2AF1^wt/S34F^*cells. Notably, *U2AF1^wt/S34F^* cells were significantly more dependent on mitochondrial respiration for their energy requirements, as evidenced by a substantial reduction in puromycin incorporation upon oligomycin treatment (**Fig. 5C-D**). Thus, the presence of the U2AF1-S34F mutation leads to metabolic remodeling, resulting in increased dependence on mitochondrial respiration and elevated global translation, which demands higher energy consumption.

**Figure 5.**
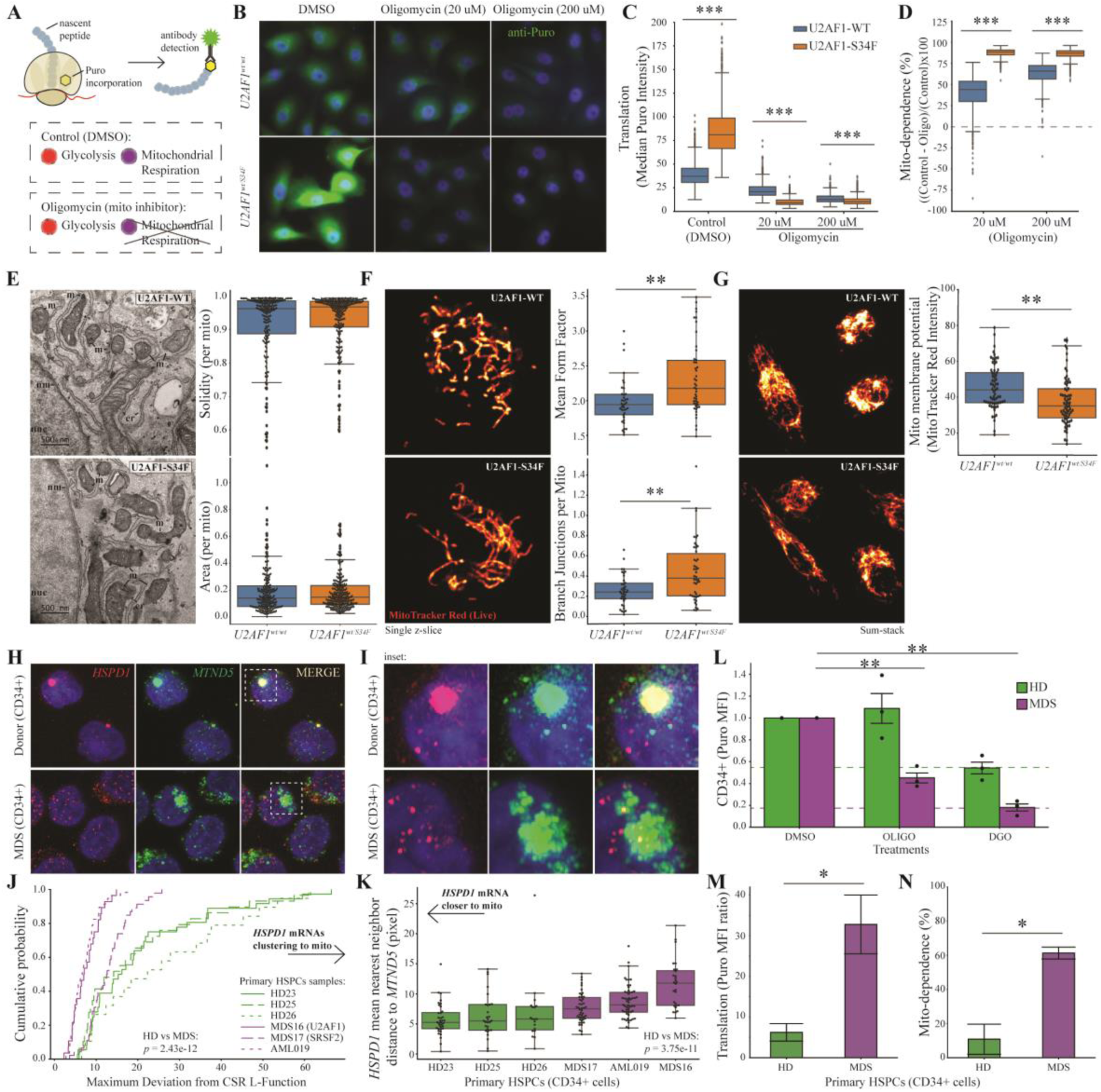
U2AF1-S34F mutant cells and primary MDS progenitors exhibit metabolic reprogramming dependent on mitochondrial function. (**A**) Schematic of protein synthesis assay. Protein translation, a proxy for intracellular energy availability, is evaluation as a function of puromycin incorporation into nascent peptides. (**B**) Representative immunofluorescence images showing puromycin incorporation in *U2AF1*^wt/wt^ and *U2AF1^wt/S34F^*cells under control conditions (DMSO) and following oligomycin-mediated inhibition of mitochondrial metabolism (20 μM and 200 μM). Puromycin incorporation is visualized by staining with an AlexaFluor488-coupled anti-puromycin antibody (green). Nuclei are DAPI stained (blue). (**C**) Box plots of median puromycin staining intensity for *U2AF1^wt/wt^* and *U2AF1*^wt/S34F^ cells in control (DMSO) and oligomycin-treated conditions (20 μM and 200 μM). Box plots display the median (line within the box) and interquartile ranges (IQR, box), and whiskers represent 1.5 times the IQR. Outliers are shown as individual points beyond the whiskers. Statistical significance was determined using the Mann-Whitney U test. \**p* < 0.05, \*\**p* < 0.01, \*\*\**p* < 0.001. (**D**) Quantification of mitochondrial dependence per cell was calculated as ((Control−Oligo)/Control)×100. The median value of the respective control sample group was used for this calculation. Cells below the dotted line indicate mitochondrial dependencies below zero, representing cells with higher puromycin staining intensity than the median value from the respective control sample. Data are presented and statistically analyzed in the same manner as in (C). (**E**) Transmission electron microscopy (TEM) images of *U2AF1^wt/wt^* and *U2AF1^wt/S34F^* cells at 6500X magnification, showing labeled mitochondria (m), endoplasmic reticulum (er), nucleus (nuc), and nuclear membrane (nm). The top box plot on the right displays the solidity per mitochondrion, a shape descriptor. The bottom box plot shows the area (in pixels) per mitochondrion. Box plots and statistical testing as in panel C, with individual mitochondria data points shown as black dots. (**F**) Super-resolution confocal images (Airyscan) of mitochondrial potential in *U2AF1^wt/wt^* and *U2AF1^wt/S34F^* cells, evaluated as a function of MitoTracker Red staining (100 nM). Regions with the highest fluorescence intensity are visualized in white. The top plot shows the mean form factor, a shape descriptor indicating mitochondrial morphology. The bottom plot shows the number of branch junctions per mitochondrion, a measure of mitochondrial connectivity. Box plots and statistical testing were performed as in panel C, with individual cell data points shown as black dots. (**G**) Sum-stack super-resolution confocal images (Airyscan) of mitochondrial membrane potential in *U2AF1^wt/wt^* and *U2AF1^wt/S34F^* cells, as a function of MitoTracker Red (100 nM) staining. The right box plot displays the median MitoTracker Red intensity. Box plots and statistical testing were performed as in panel C, with individual cell data points shown as black dots. (**H**) Representative smFISH images from primary CD34^+^ HSPCs isolated from healthy donors (HD) and MDS patients, probing for *HSPD1* (red) and *MTND5* (green) mRNAs, with nuclei stained with DAPI (blue) and inset displayed in (**I**). (**J**) CDF plot showing the maximum deviation from CSR L-function, assessing the clustering of *HSPD1* mRNAs to *MTND5* in primary CD34+ HSPCs from individual HD and MDS patient samples. Patients with splicing factor mutations are indicated on plot. Statistical comparisons between HDs and MDS groups were performed using the Mann-Whitney U test, with significance indicated by the *p*-value on the plot. (**K**) Mean nearest neighbor distance analysis of *HSPD1* mRNA relative to *MTND5* in CD34^+^ HSPCs isolated from individual primary HD and MDS patient samples. Data are presented as rank-ordered box plots. Statistical significance was tested as in panel O. (**L**) Puromycin-based FACs analysis of protein translation was performed on CD34+ cells isolated from healthy donors and MDS patients. Puromycin mean fluorescence intensity (MFI) is shown for CD34+ cells under control (DMSO), oligomycin-treated (O), and combined 2-DG and oligomycin-treated (DGO) conditions. DGO represents background fluorescence intensity. Bar graphs display the MFI with error bars representing SEM. Data represent three biological replicates (*n* = 3) for each sample group. Statistical significance was determined using a *t*-test; \**p* < 0.05, \*\**p* < 0.01, \*\*\**p* < 0.001. (**M-N**) Quantification of translational readout (Puro MFI Ratio) (**M**) and mitochondrial dependence (mito-dep) (**N**) in CD34+ HSPCs from HDs and MDS patients.

This metabolic remodeling coincides with changes in the spatial network and function of mitochondria. TEM ultrastructure analysis revealed no significant differences in individual mitochondrial shape (solidity) or size (area) between *U2AF1^wt/wt^* and *U2AF1^wt/S34F^* cells (**Fig. 5E**). However, *U2AF1^wt/S34F^*cells exhibited significantly altered mitochondrial morphology and decreased membrane potential, as quantified with MitoTracker live-staining (**Fig. 5F-G**). These findings indicate that the U2AF1-S34F mutation leads to alterations in mitochondrial network and function, despite no changes being observed at the ultrastructural level.

Finally, we directly investigated mitochondrial function in the context of diseases driven by somatic splicing factor mutations. Myelodysplastic syndromes (MDS), clonal stem cell disorders characterized by bone marrow failure and mitochondrial dysfunction, are notably associated with a high frequency of mutations in splicing factors involved in 3’ splice site selection.^17,19,20^ As a functional class, splicing factors, including U2AF1, SRSF2, ZRSR2, and SF3B1, represent the most commonly mutated pathway in MDS. Therefore, we aimed to determine whether the phenotypes observed in our tissue culture model of the U2AF1-S34F mutation are also present in bone marrow (BM) hematopoietic stem and progenitor cells (HSPCs) isolated from MDS and acute myeloid leukemia (AML) patients, including those with splicing factor mutations (**Table S5)**. First, we performed two-color smFISH using probes for *HSPD1* mRNA and the mitochondrial marker *MTND5* in BM CD34+ HSPCs from age-matched healthy donors as compared to MDS/AML patients. Notably, two of the MDS patients included in this analysis contain oncogenic splicing factor mutations. Strikingly, *HSPD1* mRNA and mitochondria showed a strong spatial co-localization in healthy donor HSPCs (**Fig. 5H-I**). Approximately 40% of these cells displayed large asymmetric assemblies of mitochondria and *HSPD1*, reflected in the long tail of Ripley’s L-function analysis (**Fig. 5J**). In contrast, patient-derived CD34+ cells exhibited disrupted mRNA targeting to mitochondria across genotypes (**Fig. 5J-K**), with the U2AF1-S34F patient sample (MDS16) showing the most pronounced disruption. In summary, healthy donor HSPCs display a degree of mitochondrial mRNA localization not previously observed in human tissue culture cells, and this co-localization is severely curtailed in MDS HSPCs.

These data strongly suggested that MDS CD34^+^ HSPCs would also exhibit differences in mitochondrial respiration relative to healthy donors. Using a puromycin-based FACs analysis of protein translation, we evaluated protein synthesis in CD34+ HSPCs in response to inhibition of mitochondrial ATP synthesis using oligomycin (**Fig. 5L**). The basal level of protein translation was approximately 5-fold higher in MDS HSPCs as compared to healthy donor HSPCs (**Fig. 5M**). Moreover, these HSPCs exhibited significantly different dependence on mitochondrial respiration. Consistent with previous studies showing the reliance of HSPCs on glycolysis in the hypoxic BM niche^47^, healthy donor HSPCs were not dependent on mitochondrial respiration—treatment with oligomycin did not significantly alter protein translation (**Fig. 5L**). In contrast, MDS HSPCs were markedly dependent on mitochondrial respiration. Oligomycin-mediated inhibition of OXPHOS significantly decreased puromycin incorporation by 2-fold, demonstrating a mitochondrial dependence of 63% (**Fig. 5L,N**). Thus, similar to the *U2AF1^wt/S34F^* cell line, primary CD34+ HSPCs from MDS patients exhibit increased basal translation, altered mRNA targeting to mitochondria, and metabolic remodeling with an increased dependency on mitochondrial respiration. Importantly, these findings provide a genotype-phenotype connection between a somatic driver mutation (U2AF1-S34F) and mitochondrial dysfunction in HSPCs.

## Discussion

We propose that the U2AF heterodimer plays a non-canonical role in the translation, trafficking, and localization of nuclear-transcribed mRNAs which code for mitochondrial proteins. A single heterozygous missense mutation (S34F) in the ZnF domain of U2AF1 results in multi-faceted translational and mitochondrial dysfunction, recapitulating defining bone marrow phenotypes in MDS. More generally, these data provide a novel mechanism for how nuclear RBPs might enable coupling between the nucleus and mitochondria.

Insights from two existing molecular paradigms— ER targeting and the exon junction complex (EJC)—may offer clues into the mechanisms underlying U2AF’s non-canonical role. In ER targeting, the signal recognition particle (SRP) binds the N-terminal signal sequence peptide and stalls translation until the nascent peptide docks at the translocon.^48^ Similarly, while mitochondrial proteins possess N-terminal targeting sequences, no functional analog of the SRP has been identified for mitochondria.^49^ We have shown that U2AF represses translation initiation and/or elongation in this MTS region and also interacts with mitochondrial proteins.^50^ This activity is altered by mutations either in the U2AF binding site on the mRNA or in the U2AF protein itself and depends on the proximity of the binding site to the start codon. Interestingly, the nascent peptide is still in the ribosome exit tunnel for positions of ∼ 90 – 120 nt^49^, which is similar to the region where U2AF binds. We propose that U2AF functions as a kinetic block to nascent peptide synthesis, facilitating the recruitment of chaperones or other RBPs necessary for molecular targeting to the mitochondria. For example, the protein CLUH has been demonstrated to interact with RNA and have a role in mitochondria biogenesis, but CLUH has no known binding RNA motif.^51^ Although our data do not distinguish between post-translational and co-translational protein import pathways, emerging evidence indicates that RNA localization works in concert with translation, supporting a direct role for U2AF in this process.^29,30^ For the transcripts we examined in depth, mutations in the U2AF1 RNA binding domain lead to concurrent changes in translation, mRNA localization, and protein targeting to mitochondria.

The EJC, which is also connected to RNA splicing, is deposited in the nucleus and aids in mRNA trafficking, translation, and quality control.^52^ It is subsequently removed during the pioneer round of translation. These roles are analogous to those we propose for U2AF. However, there are key differences between the two. While the EJC is deposited ∼ 20 nt upstream of exon-exon splice junctions with no underlying sequence specificity, U2AF binding sites are located in coding regions with a well-defined specificity, including on intronless genes. Therefore, we propose that U2AF’s role in translation and trafficking is independent of splicing, which is consistent with our in vitro findings. Moreover, recent data indicates the EJC is also associated with stable mRNPs containing stalled ribosomes^53^, analogous to what we observe for U2AF. Yet, when collided ribosome fractions were isolated after sucrose density fractionation, EJC components were not reported, but splicing factors, including components of the SF3 complex and U2AF were identified.^54,55^

This non-canonical function of U2AF is particularly apparent in primary CD34+ HSPCs, which are profoundly affected by somatic mutations in splicing factors. In particular, we see substantial co-localization between the *HSPD1* mRNA and mitochondria in healthy donor cells which is abrogated in MDS HSPCs. Additionally, we observed a metabolic re-modeling with an increased reliance on mitochondrial energy production in MDS HSPCs. Mitochondrial respiration in hematopoietic stem cells is crucial for balancing cell fate decisions between self-renewal and differentiation.^56,57^ Under normal conditions, HSPCs maintain quiescence and low energy demands, primarily relying on glycolysis with minimal mitochondrial activity.^47^ In MDS, we propose that the U2AF1-S34F mutation leads to altered mRNA translation and targeting to mitochondria, disrupting important post-transcriptional gene regulation and subsequently mitochondrial metabolism in HSPCs.

### Limitations of the study

While our study offers a new framework for understanding U2AF1 biology, particularly its role in mitochondrial-associated regulation in both normal and disease contexts, it also highlights significant gaps in our mechanistic understanding. Specifically, the precise mechanisms by which U2AF impedes translation and concurrently impact the localization of mRNAs to mitochondria remain to be elucidated. Additionally, the processes governing the trafficking of U2AF between the cytoplasm and nucleus, including the timing and regulation of its nuclear import, are still unclear. Although we speculate that U2AF binds co-transcriptionally and forms part of an export-competent mRNP, future research is needed to uncover these mechanistic details.

Finally, while the phenotype of disrupted mitochondrial homeostasis due to the U2AF1 S34F mutation is clear, it is difficult to completely rule out splicing changes—which unavoidably occur in this mutant background—as the underlying cause. However, these splicing changes are variable and modest for this mutation, with one comprehensive analysis reporting only 16 splicing changes which could be definitively linked to the mutation, none of which with known connections to mitochondria biology.^58^ Future efforts should aim to develop separation-of-function mutants for U2AF and other SR proteins to distinguish between splicing-dependent and splicing-independent effects.

## Acknowledgments

We thank members of the Larson laboratory and the NIH Translation SuperGroup for helpful discussions. We thank the many support staff at the NIH core facilities, including Guofeng Zhang (Electron Microscopy Unit), Tatiana Karpova (Optical Microscopy Core), Kathy McKinnon (CCR Flow Cytometry Core), the Frederick Mass Spectrometry Unit, and the NIH High-Performance Computing Group (Biowulf). We thank the USF-TGH Precision Medicine Biorepository for supplying the human bone marrow samples. We also thank Jonthan Yewdell for generously providing the Anti-Puromycin AlexaFluor647 PMY-2A4 antibody. We acknowledge the use of OpenAI’s GPT for assistance with coding.

## Author contributions

Conceptualization: GRG, MP, DRL

Methodology: GRG, MP, BD, BW, DS, DRL

Investigation: GRG, MP, BD, JM, KM, IP

Visualization: GRG, MP, DRL

Funding acquisition: DRL

Project administration: GRG, KM, DRL

Supervision: DRL

Writing – original draft: GRG, MP, BD, DRL

Writing – review & editing: GRG, MP, JM, BD, BW, NT, KM, BS, DMS, GR, DRL

## Supplemental information

Document S1. Figures S1–S4 and Tables S3 and S5

Table S1. Excel file containing quantitative mass spectrometry analysis of U2AF1-interacting proteins in the cytoplasmic and ER/nuclear fraction data too large to fit in a PDF, related to Figure 1

Table S2. Excel file containing U2AF1-WT cytoplasmic RNA target peaks identified by wavClusteR analysis on PAR-CLIP data too large to fit in a PDF, related to Figure 2

Table S4. Excel file containing U2AF1-S34F cytoplasmic RNA target peaks identified by wavClusteR analysis on PAR-CLIP data too large to fit in a PDF, related to Figure 2

## STAR Methods

## RESOURCE AVAILABILITY

### Lead contact

Further information and requests for resources and reagents should be directed to and will be fulfilled by the lead contact, Daniel R. Larson.

### Materials availability

Plasmids generated in this study have been deposited to Addgene (in progress, will list cat # upon acceptance)

### Data and code availability

Ribosome-sequencing data have been deposited at GEO and are publicly available as of the date of publication. Mass spectrometry data have been deposited at MassIVE and are publicly available as of the date of publication. Accession numbers are listed in the key resources table. Original western blot images have been deposited at Mendeley and are publicly available as of the date of publication. The DOI is listed in the key resources table. Microscopy data reported in this paper will be shared by the lead contact upon request.

Any additional information required to reanalyze the data reported in this paper is available from the lead contact upon request.

## METHOD DETAILS

### U2AF1-3XFLAG cell line

We used CRISPR/Cas9 to tag endogenous U2AF1 with a 3XFLAG epitope in Human Bronchial Epithelial Cells (HBECs) cultured in Keratinocyte SFM media (Gibco, 17005042). The U2AF1-3XFLAG donor plasmid, derived from the base plasmid pUC19, has approximately 800 bp flanking homology arms. Amplification of these homology arms used the following primers: Forward: TGTTGGGCTGCAGGTTGGACAAGC and Reverse: CTGGGTGACTCCCACCTAGAGC. The left homology arm spanned from chr21:43,093,105 to 43,093,960, while the right homology arm extended from chr21:43,092,220 to 43,093,098. A gene block facilitated the incorporation of the 3XFLAG epitope, T2A sequence, and a blasticidin resistance marker into the donor plasmid.

To target the C-terminus of U2AF1, we cloned the sgRNA sequence (GACATAAGGTAAAAATGGCA) into the PX458 plasmid. HBECs underwent electroporation with both the CRISPR PX458 plasmid and the U2AF1-3XFLAG donor plasmid following the manufacturer’s instructions, using the following settings: voltage = 1400, width = 20, pulses = 2 (ThermoFisher, N1025). Seven days post-electroporation, cells underwent antibiotic selection with 10 μg/mL of blasticidin for 12 days, with media changes conducted daily for the initial 5 days, followed by changes every two days thereafter. Fresh antibiotics were added to the media for each media exchange during the selection period. Integration of the 3XFLAG epitope into the U2AF1 gene locus was confirmed by PCR and sanger sequencing on cDNA using primers designed to detect both the wild-type (167 bp band) and integrated transcripts (740 bp band) using the following primers: Forward: TCTAGAGACCGTGGTCGTGG, and Reverse: CTGGTTTGGTCAATCAACTACAACAC.

### U2AF1-3XFLAG IP-mass spectrometry

Human Bronchial Epithelial Cells (HBECs) expressing endogenously tagged U2AF1-3XFLAG underwent subcellular fractionation as previously described ^59^. In addition, a parental HBEC line lacking the endogenous U2AF1 tag was prepared to serve as a control. This control was critical for identifying nonspecific interactions, ensuring that enriched proteins were attributable to the specific interactions of U2AF1-3XFLAG. The experiment was conducted with three biological replicates per sample group, each consisting of approximately 20 million cells. Briefly, cells washed with PBS at room temperature were detached using trypsin, collected in ice-cold PBS, and transferred to pre-chilled 15 ml conical tubes to inhibit degradation. After centrifugation at 500 x g for 5 minutes at 4°C, the supernatant was removed, and the cell pellet weighed. The pellet was then washed with ice-cold 1X PBS and resuspended in hypotonic lysis buffer (HLB) at a ratio of 1 ml per 75 mg of cells, with a 1X final concentration of protease inhibitor (PI) cocktail and phosphatase inhibitor (PhI) solution. Following a 10-minute incubation on ice with intermittent vortexing, the suspension was centrifuged at 800 x g for 10 minutes to collect the cytoplasmic fraction. The nuclear pellet underwent four washes with HLB before being resuspended in nuclear lysis buffer (NLB) at half the initial HLB volume, with PI and PhI added. Nuclei were then sonicated ten times, alternating between 30 seconds of sonication and 30 seconds of cooling. Subsequent centrifugation at 18,000 x g for 15 minutes at 4°C clarified the nuclear extracts.

For immunoprecipitation (IP), 30 µL of magnetic beads (specific for FLAG-tag affinity) were prepared following the manufacturer’s instructions (Sigma, M8823), washed with NLB, and incubated with either 1 mL of cytoplasmic extract or 500 µL of nuclear extract overnight at 4°C with rotation to ensure thorough binding.

#### On-bead digestion and TMTpro labeling

Afterward, beads were washed three times to remove nonspecifically bound components. Samples were digested on bead by treatment with 50µL each of lysis buffer, reducing buffer, and alkylation buffer provided with the EasyPep kit (Thermo, A40006) then incubated at 25°C 1.5hrs in the dark. Samples were treated with 20µL of 100ng/µL trypsin/LysC and incubated at 37°C overnight for 20hrs with shaking at 1000rpm. Removed the supernatant off the beads to a clean tube then treated with 50µL of 5µg/µL TMTpro in 100%ACN and incubated at 25°C for 2hrs. Samples were then quenched with 50µL of 5% hydroxylamine, 20%FA and incubated for 20 minutes before combining samples within each plex. Each plex was cleaned using EasyPep mini spin columns (Thermo, A40006) and drying eluted peptides by speed-vac.

#### LC-MS/MS analysis

Each plex was loaded twice onto a Dionex U3000 RSLC in front of a Orbitrap Eclipse (Thermo) equipped with an EasySpray ion source Solvent A consisted of 0.1%FA in water and Solvent B consisted of 0.1%FA in 80%ACN. Loading pump consisted of Solvent A and was operated at 7 μL/min for the first 6 minutes of the run then dropped to 2 μL/min when the valve was switched to bring the trap column (Acclaim™ PepMap™ 100 C18 HPLC Column, 3μm, 75μm I.D., 2cm, PN 164535) in-line with the analytical column EasySpray C18 HPLC Column, 2μm, 75μm I.D., 25cm, PN ES902). The gradient pump was operated at a flow rate of 300nL/min and each run used a linear LC gradient of 5-7%B for 1min, 7-30%B for 133min, 30-50%B for 35min, 50-95%B for 4min, holding at 95%B for 7min, then re-equilibration of analytical column at 5%B for 17min. All MS injections employed the TopSpeed method with three FAIMS compensation voltages (CVs) and a 1 second cycle time for each CV (3 second cycle time total) that consisted of the following: Spray voltage was 2200V and ion transfer temperature of 300 ⁰C. MS1 scans were acquired in the Orbitrap with resolution of 120,000, AGC of 4e5 ions, and max injection time of 50ms, mass range of 350-1600 m/z; MS2 scans were acquired in the Orbitrap using TurboTMT method with resolution of 15,000, AGC of 1.25e5, max injection time of 22ms, HCD energy of 38%, isolation width of 0.4Da, intensity threshold of 2.5e4 and charges 2-6 for MS2 selection. Advanced Peak Determination, Monoisotopic Precursor selection (MIPS), and EASY-IC for internal calibration were enabled and dynamic exclusion was set to a count of 1 for 15sec. The only difference in the methods was the CVs used, one method used CVs of −45, −60, −75 and the second used CVs of −50, −65, −80.

#### Database search and post-processing analysis

Raw files from each plex were batched together as fractions and searched with Proteome Discoverer 2.4 using the Sequest node. Data was searched against the Uniprot Human database from Feb 2020 using a full tryptic digest, 2 max missed cleavages, minimum peptide length of 6 amino acids and maximum peptide length of 40 amino acids, an MS1 mass tolerance of 10 ppm and MS2 mass tolerance of 0.02 Da. Variable modifications of oxidation on methionine (+15.995 Da) and TMTpro (+304.207) on lysine and peptide N-terminus as well as fixed carbamidomethyl on cysteine (+57.021). Percolator was used for FDR analysis and TMTpro reporter ions were quantified using the Reporter Ion Quantifier node and normalized on total peptide intensity of each channel. Proteins were filtered using a FDR cutoff of 1% and have quantitative values in at least 4 samples to be included in the final list. To further elucidate the biological significance of the identified protein interactions, we performed Gene Set Enrichment Analysis (GSEA) using the Broad Institute Molecular Signatures Database. We utilized the c2.cp.kegg.v2023.1.Hs.symbols.gmt gene set collection and analysis was conducted with default parameters, employing the ‘Signal2Noise’ metric, defined as (enrichment_IP_-enrichment_control_)/(std dev_IP_+std_dev_control_).

### Western blot validation of IP-Mass Spec

For sample preparation, anti-FLAG M2 magnetic beads (Sigma, M8823) were used to capture U2AF1-3XFLAG interacting proteins according to the manufacturer’s instructions. Briefly, magnetic beads were washed and equilibrated with HLB or NLB buffers for cytosolic and ER-nuclear fractions, respectively. A 50 µL aliquot was removed from each sample to serve as input before adding the magnetic beads. Then, 30 µL of magnetic bead solution was added to 500 µL of thawed lysate (previously flash frozen and stored at −80°C). The mixture was incubated at 4°C overnight with end-over-end rotation. After incubation, the samples were washed three times with the appropriate buffer and eluted with 3X FLAG peptide (Sigma, F4799) as per the manufacturer’s instructions. For the Western blot analysis, 0.1% input and immunoprecipitated samples were separated on a 4-12% Bis-Tris gel (Invitrogen, NP0322BOX) and transferred to a nitrocellulose membrane. Primary antibodies were incubated in the blocking solution overnight at 4°C with gentle rocking, using the following dilutions: FLAG at 1:1000 (Sigma, F3165), U2AF2 at 1:1000 (Santa Cruz, sc-53942), EIF3J at 1:500 (Bethyl, A301-746A-M), GIGYF2 at 1:500 (Bethyl, A303-732A), and G3BP1 at 1:2000 (Invitrogen, PA5-29455). Secondary antibodies (anti-mouse and anti-rabbit) were incubated in TBST for 1.5 hours at room temperature with gentle rocking. Chemiluminescence was used to detect proteins on the blot (Pierce ECL Western, 32106 or Thermo Fisher, 34094). All Western blot validations were conducted independently twice.

### DNA-PAINT single-molecule localization microscopy

The U2AF1-3XFLAG and parental control HBECs were fixed on coverslips (Deckgläser, 0117580) with 2.4% PFA for 15 minutes at room temperature. After three washes with PBS, the cells were treated with 0.1% Trion X-100 for 10 minutes. Subsequently, the samples were blocked with blocking buffer (2% BSA in 1x PBS) for 30 minutes. Following blocking, the cells were incubated with mouse anti-flag antibody (1:300; Sigma, F3165) and Rabbit anti-Tomm22 antibody (1:300; ThermoFisher, PA5-51804) for 1 hour at room temperature. After incubation, the cells were treated with anti-mouse IgG and anti-Rabbit IgG conjugated with docking sites overnight (Massive Photonics, Germany). The samples were then washed three times with washing buffer (Massive Photonics, Germany). Subsequently, 1 nM imager strands corresponding to different docking sites were sequentially added to the sample for two-color DNA-PAINTING super-resolution imaging. PBS was used for washing samples between imaging rounds until no blinking was observed.

#### Coverslip treatment

The No. 1.5 coverslips were first coated with PLL (Sigma) at room temperature for 10 minutes. After that, the coverslips were incubated with multicolor Tetraspeck beads (0.1 µm, 1:10^6^ dilution, Life Technologies) for 10 min and washed with PBS three times after incubation as fiducial marker.

#### Imaging

Imaging was performed on a custom build microscope with a Nikon Ti base and a Nikon N-STORM module equipped with a 100x 1.49 NA. oil-immersion lens. Images were acquired with a sCMOS CAMERA (Prime 95B, Teledyne Photometrics) with inclined illumination. 637 nm laser (Coherent, USA) was used as excitation wavelength. Typically, exposure time was set at 200 ms per frame for DNA-PAINTING experiments. The raw data were analyzed and rendered using the Picasso software package ^60^. The fluorescence beads were used for drift correction as well as sequential image registration. PolynomialTransformation2D algorithm was applied for different imaging rounds registration. The displayed images were minimally processed using a bandpass filter to selectively enhance the contrast of the DNA-PAINT signals while minimizing background noise.

### U2AF PAR-CLIP re-analysis

We re-analyzed previously published cytoplasmic U2AF PAR-CLIP sequencing data, which is available on the NCBI Short-Read Archive (SRA) under the Gene Expression Omnibus accession number GSE126912. Sequencing reads were aligned to the human genome (hg19) using Bowtie ^61^ with the following parameters: -v 2 (allowing up to 2 mismatches), -m 10 (reporting up to 10 alignments), and --best --strata (ensuring the best alignment for each read). We used the peak-calling program wavClusteR to identify U2AF mRNA targets as described in (<u>Bioconductor - wavClusteR</u>). Peaks were filtered to include only those located in the 5’ UTR, CDS, or 3’ UTR to focus on processed mRNAs in the cytoplasm. To generate a list of U2AF targets, we selected mRNAs that were identified in at least 2 out of the 3 biological replicates or were called in at least 1 biological replicate by the wavClusteR analysis and also identified in the original published analysis. Clusters were annotated using iterative rounds of bedtools intersect, with the following hierarchy: start codon, stop codon, coding sequence, 5’ untranslated region, and 3’ untranslated region ^62^. To assess whether U2AF binds to mitochondrial mRNAs more frequently than expected by chance, we generated two sets from U2AF1 PAR-CLIP results: a binder set (U2AF1+) and a non-binder set (U2AF1-). We then performed a Fisher’s exact test using the MitoCarta 3.0 list of human mitochondrial-associated mRNAs from the Broad Institute ^63^ to determine the significance of the intersection between these sets and the mitochondrial functional annotation. Higher order contingency testing was executed with the CTI_TEST function in IDL, which tests the indendependence using a chi-sequre goodness-of-fit test and computes the expected amount of random overlap. The inputs into this contingency table test are the overlapping and non-overlapping mRNA targets from our data and published sources encompassing: U2AF targets (2736, our analysis), mitochondrial targets (1136, see above), EIF3 targets (479, ^33^, and FMRP (1589 targets, log 2-fold enrichment >= 0.6, ^32^. For example, there are 212 hits which are common between U2AF and EIF3 par-clip datasets, and the expected level for random overlap is 56, so the ratio of observed/expected is 3.8.

#### Functional enrichment analysis

To identify enriched functional terms and pathways, we used the Enrichr online resource (<u>Enrichr (maayanlab.cloud)</u>). The list of U2AF target mRNAs, derived from our PAR-CLIP analysis, was input into Enrichr to perform the enrichment analysis. This allowed us to uncover significant biological processes, molecular functions, and cellular components associated with U2AF targets.

#### Mitochondria pathway analysis

For mitochondria specific pathway analysis, we used the Human MitoCarta3.0 database to categorize the U2AF target mRNAs into various mitochondrial pathways. We plotted the distribution of U2AF target mRNAs across these pathways using a pie chart to visualize how U2AF target mRNAs are distributed among different mitochondrial functions.

#### Meta-transcript profile analysis

We created a metagene profile using the deeptools (v.3.5.0) suite of python scripts. First, we computed the matrix using computeMatrix scale-regions ^64^. The parameters used included an upstream and downstream region of 500 bp, a bin size of 5 bp, and skipping zero values. To normalize the signal across genes, we calculated the sum of signals for each gene, normalized each value by the gene’s sum, and applied a scaling factor of 10. This normalization step ensured that the signals were comparable across different genes. Finally, we generated the heatmap using the plotHeatmap tool with the following parameters: color list (black, chartreuse, white), plot type (fill), missing data color (grey), heatmap height (15), zMin (0), and zMax (0.1), and generating standard error (SE) bands for visualization.

#### Comparative analysis of U2AF1-WT and U2AF1-S34F binding

To compare U2AF1 binding between the WT and S34F genotypes, we first identified the union of called peaks for both U2AF1-WT and U2AF1-S34F. For each genotype, we extracted reads from both the WT and S34F peaks and used the deeptools bigwigCompare tool to calculate the log2 fold change. The computeMatrix command was utilized to generate a matrix, centering on the peaks with a bin size of 5, and extending 100 bases upstream and downstream from the peak center. We skipped zeros and the output matrix was then normalized using a custom Python script to ensure accurate comparisons (see *Meta-transcript Profile Analysis* methods). The plotProfile command was employed to generate profiles of the log2 fold change. Key parameters included using a mean average type and generating standard error (SE) bands for visualization.

### RNA Bind-N-Seq (RBNS)

Our protocol for RNA Bind-N-Seq closely follows the protocol developed by the Burge Lab ^65^. The random RNA library was prepared by annealing the T7 oligo to the RBNS template, which is a single-stranded DNA oligo from IDT (sequences are listed in table below). We then performed *in vitro* transcription for 4 hours using the NEB HiScribe T7 RNA synthesis kit. After RNA synthesis, we degraded the DNA template by DNaseI digestion for 15 minutes. RNA was purified using the NEB Monarch RNA cleanup kit and stored at −80C. We confirmed that RNA was the correct length by tapestation analysis (Agilent).

U2AF (WT and S34F at 0 nM, 20 nM, 80 nM, or 320 nM) was incubated with 500 nM of random RNA library in U2AF binding buffer without tRNA (25 mM Tris pH 7.0, 150 mM NaCl, 0.5 mM DTT, 10% glycerol, 0.05% Tween 20) at 4C for 30 minutes with constant mixing on a rotisserie tube rotator. We performed each binding reaction in duplicate. Final volume was 250 uL per reaction. RNA in complex with U2AF was isolated by U2AF1-FLAG IP using anti-FLAG M2 magnetic beads (Sigma Aldrich). Beads were washed 4x in RBNS wash buffer (25 mM Tris pH 7.5, 150 mM KCl, 60 ug/mL BSA, 0.01% Tween 20) and then resuspended in 200 uL of binding buffer. We added 20 uL of washed beads to each RBNS reaction and incubated for 30 minutes at 4C. Bound RNA was pelleted with magnetic beads and incubated in 1000 uL RBNS wash buffer for 15 minutes at 4C. Next, the RNA was pelleted and resuspended in elute buffer (25 mM HEPES pH 7.3, 100 mM NaCl, 200 ug/ml 3XFLAG (sigma), 1 mM DTT). U2AF was removed from RNA by phenol-chloroform extraction and ethanol precipitation. Adding 4 uL glycogen to each ethanol precipitation significantly improved the ability to visualize such a small pellet. After ethanol precipitation, the quality of RNA was assessed by tape station.

Isolated RNA was converted to cDNA using the ImProm-II reverse transcription system (Promega) with the RT primer (see table). We also performed reverse transcription on the random RNA library not subjected to U2AF or IP (input sample). We performed PCR on each sample using NEBNext Multiplex Oligos for Illumina (NEB). We assessed PCR product by agarose gel and chose the number of amplification cycles to avoid side products (0 nM-15 cycles, 20 to 320 nM-11 cycles, input-9 cycles). Finally, DNA bands corresponding to about 160 bp were purified from 8% Native PAGE and incubated in 400 uL gel extraction buffer (10 mM Tris, pH 7.0, 300 mM NaCl, 2 mM EDTA) for 1 hour at 65C. DNA was further cleaned using a DNA Clean & Concentrator Kit (Zymo). The correct DNA length (160 bp) was confirmed by tape station.

DNA was pooled into a single tube (67 fMoles per reaction) and sent for deep sequencing at the NCI sequencing facility. Sequencing was performed on a NovaSeq (Illumina) in an S2 flow cell with 2×100 read length. We sequenced to a depth of between 149 million to 450 million reads per binding reaction which allowed us to determine enrichment scores for very long U2AF binding sites. For all samples, at least 90% of reads had a quality score above Q30.

Analysis, performed using the RBNS pipeline (https://github.com/cburgelab/RBNS_pipeline), was too computationally intensive for a single computer. Therefore, the pipeline was performed on Biowulf, a linux cluster maintained by the NIH High Performance Computing center.

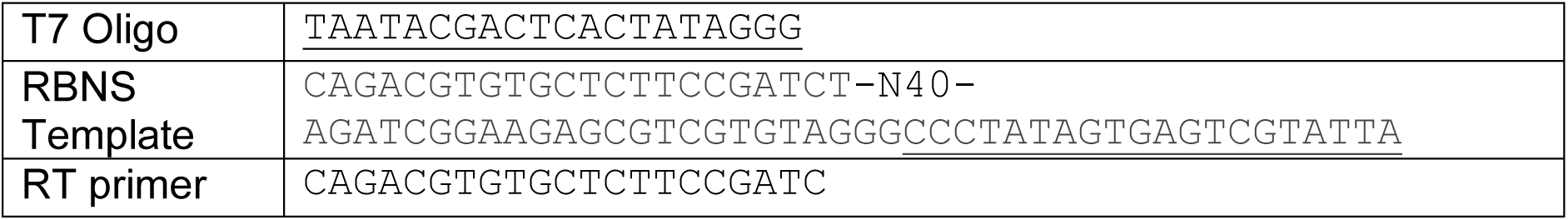

#### Predicting relative binding affinities of in vivo U2AF binding sites

We randomly selected 2.5 million reads from each U2AF concentration of the RNA Bind-N-Seq assay for analysis in the probound server (http://probound.bussemakerlab.org/) (20). Each run also included 2.5 million reads from the input fraction. We generated U2AF binding models for 6mer, 12mer, and 18mer sequences (see the accompanying paper by Donovan *et al*.).

We used the 12mer binding model from U2AF = 20 nM to assign affinities to each peak from the cytoplasmic U2AF PAR-CLIP experiment. We determined the U2AF binding affinity of every 12mer in a single PAR-CLIP peak and assigned the highest affinity value as affinity of that binding site. We did not consider the small fraction of PAR-CLIP peaks that spanned less than 12 nt.

### U2AF motif analysis on cytosolic and mitochondrial-associated mRNAs

Mitochondria-targeted and non-targeted genes are based on the annotation in MitoCarta3.0. The mitochondria-targeted gene list was used as input into Ensembl BioMart to download the protein coding sequence from GRCh38.p14. As the input to BioMart is a gene name, multiple isoforms are returned, and all isoforms are used in subsequent analyses. This list of CDS is input into the FIMO algorithm from the meme suite (https://meme-suite.org/meme/tools/fimo). The 3’ SS motif is taken directly from ^37^ and a motif search is executed with p<0.01. The complete function call is:

\# FIMO (Find Individual Motif Occurrences): Version 5.5.5

\# fimo --oc. --verbosity 1 --bgfile --nrdb----thresh 0.01 --norc --no-pgc motifs.meme mart_export_mito.txt

The output is a table of motifs and positions relative to the start of the CDS. For example, a search over 1,136 mitochondrial genes and respective isoforms at stringency of p<0.01 gives ∼60,000 motifs. A frequency distribution is created for all site positions in the CDS with binsize of 33 nt. The data is normalized by the frequency distribution of CDS lengths. Error bars are determined by bootstrap sampling from the original list of mitochondrial transciprs (n=10). The identical analysis was carried out for 5 random sets of ∼1,000 genes which are not mitochondrial targeted. Error bars are SEM for these 5 sets.

### Cross-correlation between PAR-CLIP and RIBO-Sequencing

#### Ribosome footprinting

Human Bronchial Epithelial Cells (HBECs) carrying either WT or the heterozygous S34F mutant allele of U2AF1 were cultured in Keratinocyte Serum Free media (Invitrogen) in 2 x 225cm dishes/cell line up to ∼80% confluency and treated with cycloheximide (100 μg/mL) for 10 min at 37 °C. After the incubation, cells were quickly washed with ice cold PBS and rapidly scrapped off the plate into lysis buffer, cells lysed and ribosomes treated with RNAseI as described previously ^66^. The RNAse digested lysate was layered on a 1M sucrose cushion (made in lysis buffer) and centrifuged in a Beckman TLA 100.3 rotor at 70,000 rpm for 3h at 4 °C. The ribosome protected RNA fragments associated with monosomes and disomes were recovered from the ribosomal pellet using a mirVana microRNA isolation kit (Invitrogen). Instead of radioactively end labeling the RNA fragments to size select the ribosome protected RNA fragments by SDS-PAGE as described in the original protocol Ingolia etal, we resorted to an alternate method described by ^40^. After purification of the monosome and disome RNA fragments, adaptor ligation, library preparation and size selection of the monosome and disome footprint fragments were performed as described previously ^40^. Briefly, after ligation of the 3’- and 5’-adaptors to the RNA fragments, the purified ligated fragments were reverse transcribed, and the cDNA was PCR amplified for 5 cycles with primers complementary to the adaptors at the 5’- and 3’-ends. This PCR product was then fractionated on a 3% Pippin gel and DNA fragments of 74-90 bp and 100-130 bp corresponding to the monosome and disome footprints respectively were size selected. The size selected DNA fragments were then PCR amplified for 7 cycles to generate the final library for sequencing.

#### Ribo-Seq sequencing and analysis

Sequencing libraries from Ribo-seq experiments were prepared in three biological replicates per genotype (wild-type and S34F mutant), for both the monosome and polysome fractions, for a total of 12 samples. Ribo-seq reads were sequenced on an Illumina NovaSeq 6000 instrument with SP flow cell for 101 cycles in single-end mode. Greater than 80 million reads per sample was generated.

Ribosome protected reads were first trimmed of low quality and adaptor sequence using cutadapt v.3.4 and the parameters ‘-m 15 --rc ‘, supplying a file of adapter sequence in fasta format ^67^. Trimmed reads were then filtered of rRNA reads by local alignment to a reference of human ribosomal sequence using Bowtie2 (v.2.4.4) ^68^. Trimmed, rRNA filtered reads were then aligned using Bowtie2 (v2.4.4) in local mode to a reference of transcript sequence of UCSC canonical transcripts, downloaded from the UCSC genome browser ^69^. Tracks in bigwig format were generated using deeptools v.3.5.0 ^64^. Depth normalized tracks were in counts per million (CPM) were used in browser visualizations, and non-normalized for cross-correlation analysis.

#### PAR-CLIP-Seq remapping (to transcriptome)

Data for PAR-CLIP experiments were re-processed to enable parallel comparison with Ribo-seq data in a common reference. Raw data were retrieved from the Gene Expression Omnibus (GEO) accession GSE126912 (U2AF), GSE65004 (EIF3), and GSE39686 (FMR1). Raw data for G3BP2 were retrieved from the Sequence Read Archive, under bioproject PRJNA533136. Reads were first aligned to the hg19 reference genome using Bowtie v.1.3.1 with the parameters ‘-v 2 -m 10 --all --best --strata --chunkmbs 384 --sam --no-unal’, to filter non-genomically derived reads. Aligned reads mapping to annotated transcripts were then extracted using samtools and bedtools intersect. These reads were mapped with Bowtie v.1.3.1 to the same canonical transcript reference as the Ribo-seq experiments. Alignments representing bona fide PAR-CLIP targets were filtered via confirming T to C changes using the samtools calmd command.

#### Cross-correlation analysis

Cross-correlation was executed using a direct application of time-series approaches but transformed into sequence space ^41^. Briefly, this calculation amounts to the cross co-variance between an RNA binding protein coverage map and either the monosome or disome coverage plot map. Instead of a time delay which is the coordinate in a time-series approach, the coordinate for genomic analysis is bases or nucleotide position. As described in detail in ^41^, the base sampling increases with separation and is logarithmic, which decreases noise at large separation. Importantly, the RBP coverage plot is only generated from T to C reads which are a signature of direct binding in the PAR-CLIP analysis. We found that including all PAR-CLIP reads generated qualitatively similar cross-correlations, but the dynamic range was decreased (data not shown).

### *In vitro* MCL1 reporter

#### Plasmids

Plasmid coding for hMCL1-NanoLuciferase RNA used for in vitro translation reporter assays were constructed as follows. Poly(A) mRNA isolated from Human Bronchial Epithelial Cells was reverse transcribed using oligo dT primers and ProtoScript II (New England Biolabs) as per the manufacturer’s instructions. Human MCL1 coding sequence to include the 5’-UTR was PCR amplified from the cDNA and cloned downstream of the SP6 promoter in the pSP64 Poly(A) vector (Promega Corporation). A custom synthesized DNA fragment (IDT) coding for Nano-Luciferase and the 3’-UTR of hFTL1 was then inserted in-frame downstream of MCL1 coding sequence. All cloning reactions were performed using restriction enzymes as prescribed by the manufacturer (New England Biolabs). The resulting plasmid, MCL1_Nano-Luciferase Reporter (MP127), was used to construct all subsequent mutant and sequence deletion variants of MCL1 RNA used in this study, either by site directed mutagenesis or using commercially obtained synthetic DNA fragments (IDT). The sequence of all plasmids was verified by Sanger sequencing.

#### Synthesis of RNA for in vitro translation assay

The template for in vitro transcription was PCR amplified from the plasmid MP127 (see above) using a forward primer (upstream of the Sp6 promoter) and a reverse primer (immediately downstream of the poly(A) tail), restricted with EcoRI and the purified fragment served as template for in vitro transcription. RNA was synthesized using the HiScribe SP6 RNA synthesis kit (New England Biolabs) supplemented with the 5’ mRNA cap analog and as prescribed by the manufacturer. The synthesized RNA was purified using the Monarch RNA purification kit (New England Biolabs).

#### Preparation of cytosolic extract for in vitro translation

Cultured HeLa S3 cell pellets shipped on wet ice were obtained from Cell Culture Company, Minneapolis, MN. Cells were suspended in ice cold hypotonic buffer and crude cell lysates prepared as described previously ^70^ and centrifuged at 3300g for 5 min at 4 °C. The supernatant was removed and centrifuged at 10,000g for 10 min at 4 °C. The supernatant, S10 cytoplasmic extract was aliquoted, snap frozen and stored under liquid N2 until further use.

#### Purification of U2AF from S9 insect cells

DNA fragments with the coding sequence for wild type or S34F U2AF1 with an N-terminal 3X FLAG tag and U2AF2 with an N-terminal 6X His tag were commercially synthesized (IDT) and cloned downstream of the p10 and polyhedron promoters respectively in the pFASTBAC baculovirus expression vector to create plasmids U2AF1-WT_U2AF2_Bac (MP66) and U2AF1-S34F_U2AF2_Bac (MP67) respectively. S9 cells were transduced with baculovirus packaged with the expression vectors. Briefly, a packed cell volume of 20mL of infected cells was used for purification of the WT and S34F U2AF1/U2AF2 heterodimer. Cells were lysed by sonication in lysis buffer (Tris-HCl 50 mM, pH 8.0; NP40 1%; Immidazole 50 mM; KCl 300mM and protease inhibitor) and the crude lysate centrifuged at 13,000rpm for 15 min at 4 °C in a Sorvall SS-34 rotor. The supernatant was bound to a HiTRAP His column, washed with 4 volumes of buffer (Tris-HCl 50 mM, pH 8.0; KCl 300 mM and Immidazole 50 mM) and protein eluted with 2 volumes of buffer (Tris-HCl 50mM, pH 8.0; KCl 100 mM and Immidazole 500 mM). Fractions containing the protein were pooled, dialyzed against dialysis buffer (Tris-HCl 20 mM, pH 8.0; NaCl 150 mM, and Glycerol 10%). The dialyzed protein was bound to Anti-FLAG M2 magnetic beads (equilibrated in dialysis buffer) for 2h at 4 °C, beads washed in the same buffer containing NaCL 500 mM and eluted with the same buffer containing NaCl 100 mM and 100 μg/mL 3XFLAG peptide. The eluted protein was dialyzed against Tris-HCl 20 mM, pH 8.0; NaCl 100 mM, and Glycerol 10%, aliquoted, flash frozen and stored at −80 °C.

#### In vitro translation and nano-luciferase assay

*In vitro* translation assay contained 20x translation assay buffer (Invitrogen) supplemented with Methionine (50 μM), RNAsin (0.5U); purified RNA (50 nM) and S10 cytosolic extract (50 μg protein) in a final volume of 5 μL. Reactions were incubated at 30 °C for 60 minutes and terminated by transferring the reactions on ice. Nano-Luciferase activity was assayed in 100 μL that contained 5 μL translation mix, 45 μL water and 50 μL Nano-Glo Luciferase Assay reagent (Promega), and luminescence measured in a SpectraMax ID3 (Molecular Devices) plate reader. All in vitro translation assays were performed at least two times with three replicates each. Statistical significance was assessed with a Students t-test of area under the curve.

### Polysome profile followed by Western-blot

Polysomes were isolated from parental and U2AF1-3XFLAG HBECs cultured in 15-cm dishes at approximately 80% confluency, following the method described by ^71^. Plates were incubated with fresh media for approximately 2 hours before starting the assay to ensure actively translating cells. To freeze ribosomes on the transcripts, cells were incubated in medium containing 300 µM Emetine for 3 minutes under normal culture conditions, washed with pre-warmed PBS, trypsinized, and washed again with ice-cold PBS containing 300 µM Emetine. Cells were then lysed in lysis buffer (20 mM Tris at pH 7.2, 130 mM KCl, 15 mM MgCl2, 0.5% [v/v] NP-40, 0.2 mg/mL heparin, protease inhibitors (Sigma), RNASin (Promega), 2.5 mM DTT, 0.5% deoxycholic acid, and 300 µM Emetine). The extract was clarified by centrifugation (8000g for 10 minutes at 4°C). To normalize RNA loading, triplicate A260 measurements from 1:10 dilutions of each clarified lysate were read using a NanoDrop UV-Vis spectrophotometer. Background A260 measurement from lysis buffer was subtracted from each reading. Equal RNA loads (approximately 100 µg) were layered on top of each sucrose gradient and centrifuged (41,000 rpm for 2 hours at 4°C) in an SW41 Ti rotor (Beckman). UV profiling (254 nm) of the fractions was performed using a fractionator and Triax Flow Cell software (Biocomp Instruments).

#### Protein isolation from polysome fractions

Protein isolation from polysome fractions was conducted as follows: Two volumes of 100% ethanol were added to each sample (e.g., 200 µL of ethanol to 100 µL of sample), with 500 µL of fraction added to 1 mL of 100% ethanol in a 1.7 mL tube. Samples were incubated overnight at −20°C, followed by centrifugation at maximum speed (13,000–14,000 rpm) for 25 minutes at 4°C. The resulting pellet was dried in a speed vacuum at low heat for approximately 10 minutes until no liquid remained. For SDS-PAGE, the protein pellet was resuspended in 30 µL of 2x Laemmli Sample Buffer (LSB). The resuspended pellet was incubated for 15 minutes at 95°C, vortexed briefly, and subjected to a spin at high speed for 5 minutes. The samples were either stored at −20°C or used immediately for SDS-PAGE, loading approximately 10-15 µL per lane on a 4-12% Bis-Tris gel (Invitrogen, NP0322BOX). Primary antibodies were incubated in the blocking solution overnight at 4°C with gentle rocking, using the following dilutions: RPS19 at 1:500 (Bethyl, A304-002A), FLAG at 1:1000 (Sigma, F1804), and U2AF2 at 1:1000 (Santa Cruz, sc-53942). Secondary antibodies were diluted at 1:2500 in TBST for 1 hour at room temperature with gentle rocking. SuperSignal West Femto chemiluminescence was used to detect proteins on the blot (Thermo Fisher, 34094). All experiments were conducted independently three times.

### 2-color smFISH (HBEC cells)

HBECs were grown to 70-80% confluency on 18mm #1 (1/2 thickness) cover glasses placed in 12-well plates, with cells seeded at least 36 hours before fixation to ensure one round of division (Deckgläser, 0117580). All volumes were 1 ml and staing/wash steps occurred with gentle rocking unless otherwise stated. For fixation and permeabilization, cells were rinsed three times with HANKS buffer (or PBS without calcium or magnesium) at room temperature, followed by fixation with 4% PFA in PBS for 10 minutes. After fixation, cells were washed twice for 10 minutes each with PBS and then permeabilized in 70% ethanol for 1 hour at room temperature and stored at 4°C for up to one week. Subsequently, cells were washed once for 10 minutes with 1x PBS and once for 5 minutes with wash buffer (10% formamide and 2x SSC) at room temperature.

For hybridization, a piece of parafilm was placed at the bottom of a petri dish. 50-75 µl of hybridization solution (2× SSC (v/v), 10% dextran sulfate (w/v) and 10% deionized formamide (v/v)) was placed on the parafilm, and cover glasses were placed cell-side down onto the probe mixture. Several moist Kimwipes were placed in the petri dish with the cover glasses to prevent dry out, and the dish was sealed with parafilm. Hybridization was carried out at 37°C for 4 hours. For washing and mounting, cover glasses were transferred cell-side up back to a 12-well plate. Cells were washed twice with pre-warmed wash buffer, incubating each time at 37°C for 30 minutes. Cells were then washed once with 2x SSC at room temperature and covered with foil, followed by a wash with PBS for 5 minutes at room temperature, also covered with foil. A small drop (∼25-35 µl) of mounting media with Dapi (Prolong Gold with DAPI, Thermo) was added to a glass slide, and the cover glass was gently placed cell-side down onto the mounting media minimizing bubbles. The glass slides were dried overnight in a dark place and stored at −20°C when not being imaged.

#### smFISH Imaging

Single-molecule FISH experiments were imaged on a custom microscope setup consisting of a RAMM (ASI) chassis and a Zeiss 40x, 1.4 N.A. objective. Z-stack images were acquired with a z-step of 500 nm, consisting of 17 z-planes in the stack. The emission path was filtered using both a quad bandpass filter and an automated emission filter wheel for imaging DAPI, GFP, Cy3, and Cy5 (VCGR-SPX-P01-PC, Chroma). An automatic stage (ASI) was used for imaging multiple fields of view with an ORCA-Flash4 V2 CMOS camera (Hamamatsu). The microscope was controlled using Micro-Manager software (Open Imaging).

#### smFISH Analysis

To perform the spot detection, z-stack images were first converted to maximum projections. Cell segmentation and mask creation was performed in CellProfiler 4.2, and spot detection and coordinate extraction was performed in python with the FISH-quant v2 big-FISH package ^72^. To quantitatively analyze the spatial distribution of nuclear-encoded mRNAs relative to mitochondria, we employed spatial point pattern analysis using Ripley’s K and L functions ^73,74^. Using RNA coordinates and cell boundaries, this approach allows us to determine the degree to which mRNAs are randomly distributed, clustered, or dispersed around mitochondria within the cell. Ripley’s K function measures the cumulative number of molecules found within a certain distance from each other, providing insight into the degree of clustering or dispersion relative to a random distribution. Ripley’s L function, a variance-normalized version of Ripley’s K function, removes scale dependence, offering a more intuitive measure of spatial patterns. By comparing the observed distributions to theoretical random distributions, we can assess the spatial relationships between nuclear-encoded mRNAs and mitochondria. We utilized the spatstat package in R (version 4.2) to perform the analysis. RNA coordinates and cell boundaries were extracted from smFISH images, and the spatial point patterns for nuclear-encoded mRNAs and mitochondria were defined using these coordinates. Each cell’s RNA and mitochondrial coordinates were manually filtered for quality.

For each cell, we created point pattern objects and defined the cell’s window based on its coordinates. We then computed the Cross-K function to analyze the spatial relationships between mRNAs and mitochondria. Edge correction was applied to account for boundary effects where parts of the neighborhood fall outside the observation window. To assess whether the observed spatial patterns differ from randomness, we generated a confidence envelope using 250 simulations under the complete spatial randomness (CSR) hypothesis. We used the slope of the L function as ^73^ our summary statistic to quantify the degree of clustering or dispersion. Linear regression was performed on the L function values to obtain the slope, which was used to compare spatial patterns across cells from different sample groups. We then plotted the cumulative distribution function (CDF) of the slopes and performed a Mann-Whitney U test to assess statistical significance.

To compare the distance of nuclear-encoded mRNAs from mitochondrial signals across different probes, we performed a mean nearest neighbor analysis using a custom python script. First, we read in the smFISH data files containing the coordinates for mRNA and mitochondrial spots. For each cell, we used the KDTree algorithm from the SciPy library to create a spatial index of the mitochondrial coordinates. We then queried the nearest neighbor distances from each mRNA spot to the closest mitochondrial spot within the same cell. The mean nearest neighbor distance for each cell was calculated and stored for further analysis. Statistical significance was assessed with a Mann-Whitney U test. All experiments were performed at least two independent times.

### Mito-enrichment Western

To generate a mitochondrial-enriched fraction, we used the Mitochondria Isolation Kit for Cultured Cells (Thermo Fisher, 89874) following the manufacturer’s instructions. Briefly, we employed the reagent-based method (Option A) and differential centrifugation to separate the cytosolic and mitochondrial fractions. For a more purified mitochondrial fraction, we modified step 7 by centrifuging at 3,000 x g for 15 minutes. For SDS-PAGE, 800 μl of cytosolic lysate was mixed with 200 μl of 5X Laemmli sample buffer, and the mitochondrial pellet was resuspended in 100 μl of 5X Laemmli sample buffer. The samples were then incubated for 5 minutes at 95°C, vortexed briefly, and centrifuged at high speed for 5 minutes.

For the Western blot analysis, 2.5 μl of the cytoplasmic fraction and 5 μl of the mitochondrial fraction were loaded onto a 4-12% Bis-Tris gel (Invitrogen, NP0322BOX) and run using MES buffer at 100V for 1 hour. The protein was then transferred to a nitrocellulose membrane using the Trans-Blot Turbo Transfer kit (Bio-Rad, 170-4270) with standard transfer settings (25V, 1.0A, 30 minutes). The membrane was then blocked with 5% milk for 1 hour at room temperature. Primary antibodies were incubated in the blocking solution overnight at 4°C with gentle rocking, using the following dilutions: Beta actin at 1:10,000 (Sigma, A2228), HSPD1 at 1:12,500 (Sigma, HPA050025), and TOMM22 at 1:1000 (Thermo Fisher, PA5-51804). Secondary antibodies were incubated in TBST for 1.5 hours at room temperature with gentle rocking.. Chemiluminescence was used to detect proteins on the blot (Pierce ECL Western, 32106), and band intensity was quantified using ImageJ. All experiments were conducted independently three times.

### Puromycin-based protein translation imaging assay

As puromycin, an aminoacyl-tRNA analog that binds to the ribosome A-site, is covalently incorporated onto the 3′ end of the nascent polypeptides, detection of puromycylated polypeptides with an anti-puromycin antibody is a measure or protein translation ^45,46^. To monitor protein translation, approximately 40,000 cells (HBECs) were seeded onto an 18 mm cover glass (Deckgläser, 0117580) in a 12-well plate 2 days prior to drug treatment. To minimize debris, both the cells and media were filtered before seeding. At the time of drug treatment, cells were between 50-70% confluent. Approximately 1-2 hours before adding inhibitors, spent media was removed and replaced with 400 µL of fresh media to ensure active translation. To assess the impact of mitochondrial energy production on translation, oligomycin was used at final concentrations of and cycloheximide were 20 µM and 200 µM (using 400 µL of a 2X concentration of Oligomycin or DMSO). As a control, cytosolic protein synthesis was inhibited by adding cycloheximide at a concentration of 100 µg/mL (from a 2x concentrated stock). Following the addition of inhibitors and controls, cells were incubated at 37°C in the dark for 15 minutes. Notably, oligomycin was used within two weeks of reconstitution with single-use aliquots stored at −20°C. Prior to use, oligomycin was thawed and sonicated for 1 minute on high to prevent precipitation and then used immediately. After the initial 15-minute incubation with the inhibitors, puromycin was added to a final concentration of 10 µg/mL (using 800 µL of a 2X stock solution), resulting in a total volume of 1600 µL. Cells were then incubated for an additional 30 minutes in the dark. Following incubation, cells were fixed with 4% paraformaldehyde (PFA) in PBS for 15 minutes, followed by three 10-minute washes with PBS. Following fixation, cells were permeabilized in 0.1% Triton X-100 in PBS for 15 minutes at room temperature, washed three times with PBS, and blocked with 5% normal calf serum in PBS for 1.5 hours at room temperature. Subsequently, cells were incubated with Alexa Fluor 488-conjugated anti-Puromycin antibody (Biolegend, 381506) diluted 1:1000 in 5% normal calf serum at 4°C overnight in the dark. After the overnight incubation, cells were washed three times with PBS. Finally, cover glasses were mounted onto glass slides using ProLong Gold with DAPI (Invitrogen) and allowed to cure overnight at room temperature.

#### Imaging protein translation

Protein translation in the absence or presence of metabolic inhibitors were imaged on a custom microscope setup consisting of a RAMM (ASI) chassis and a Zeiss 40x, 1.4 N.A. objective. Z-stack images were acquired with a z-step of 500 nm, consisting of 17 z-planes in the stack. The emission path was filtered using both a quad bandpass filter and an automated emission filter wheel for imaging DAPI, GFP, Cy3, and Cy5 (VCGR-SPX-P01-PC, Chroma). An automatic stage (ASI) was used for imaging multiple fields of view with an ORCA-Flash4 V2 CMOS camera (Hamamatsu). The microscope was controlled using Micro-Manager software (Open Imaging).

#### Quantitative analysis of translation

To quantify anti-puromycin staining intensity, acquired z-stack images were converted into a sum-stack to aggregate the fluorescence signal across all planes. Next, CellProfiler software was used to segment the cells and measure the whole-cell intensity of the anti-Puromycin signal ^75^. Statistical differences in the median intensity values between sample groups in 3 independent experiments were evaluated using a Mann-Whitney U test.

### MitoTracker Red staining

Two days prior to imaging, approximately 40,000 cells were seeded onto a Lab-Tek 2 chamber glass slide (Thermo Fisher, 177380). To minimize debris, cells were filtered before loading into the chamber. For staining, cells were treated with pre-warmed complete KSFM media (Gibco, 10724-011) containing 175 nM MitoTracker Red CMXRos (Invitrogen, 2351994) and incubated for 30 minutes at 37 °C in a light-protected incubator. Following the incubation, the stain was removed and fresh media was added. The chamber was then immediately placed on a temperature- and CO_2_-controlled stage of a confocal microscope (protected from light). Imaging commenced 45 minutes after positioning the chamber on the stage to allow cells to acclimate to the microscope stage environment.

#### MitoTracker Red imaging

Confocal images were collected on an Airyscan Zeiss LSM880 confocal microscope using the Airyscan module to enhance resolution in XYZ (120 nm x 120 nm x 380 nm) beyond the diffraction limit at 16-bit image depth. Cells were imaged live in an environmental chamber with controlled CO2 and heating, and Definitive Focus was turned on. Z-stack images were acquired using an Alpha P-Apo 100X/1.46 NA objective with a z-step of 500 nm, consisting of 12-14 z-planes in the stack. The pixel size was set to 135 nm, and the pixel dwell time was 3.35 µs. Images were acquired in the Red channel (Ex 594 nm, 0.75% laser power; Em LP 605 nm). Data collection was controlled using ZEN software on a Windows 10-based system.

#### MitoTracker Red network structure analysis

For quantitative mitochondrial network structural analysis, we used the ‘Mitochondria Analyzer’ (Version 2.1.0) ImageJ plugin, which enables semi-automated image-based analysis of fluorescently labeled mitochondria acquired through confocal microscopy, in both 2D and 3D ^76^. We conducted our analysis on high-quality single z-slice images of mitochondria. The 2D Optimize Threshold command was used to identify the appropriate settings on test samples from each set acquired under the same imaging conditions. These optimal settings were then used for the 2D Threshold batch command on the acquired images. The Analysis command was applied to the thresholded images to quantify the morphological and/or network characteristics of the mitochondrial objects, both on a per-cell and per-mito basis. For per-cell analysis, the network characteristics were normalized to both the mitochondrial count and total volume/area, providing insights into the state of the mitochondrial networks, such as shifts to a fragmented or more filamentous state. Statistical analysis was performed using the Mann-Whitney U test to assess significance. The experiments were conducted independently two times to ensure reproducibility and reliability of the results.

#### MitoTracker Red intensity analysis

To quantify the intensity of the MitoTracker Red stain, the following steps were performed. First, the acquired z-stack images were converted into a sum-stack to aggregate the fluorescence signal across all planes. Next, CellProfiler software was used to segment the cells and measure the whole-cell intensity of the MitoTracker Red stain.

Statistical differences between the median intensity values between sample groups in 3 independent experiments was evaluated using the Mann-Whitney U test.

### TEM imaging analysis

HBEC U2AF1-WT and U2AF1-S34F cells were fixed in a mixture of 2.5% glutaraldehyde and 2% formaldehyde in sodium cacodylate buffer at room temperature for 30 minutes followed by rinse with 0.1M cacodylate buffer for 3 times each for 5 minutes and post-fixed for 30 minutes in a reduced osmium solution containing 2% osmium tetroxide and 1.5% potassium ferrocyanide in 0.15 mM sodium cacodylate buffer (pH 7.4). Then the cells were incubated with a 1% thiocarbohydrazide (TCH) solution in ddH2O for 20 minutes at room temperature (RT) following being rinsed with double-distilled water (ddH2O) for 3 times each for 5 minutes. The cells were fixed with 2% osmium tetroxide in ddH2O for 30 min at room temperature, and then incubated with 1% uranyl acetate (aqueous) overnight at 4oC. After being washed with ddH2O, the samples were subjected to en bloc Walton’s lead aspartate staining as described and placed in the oven for 30 min at 60 degrees. After another 3 rinses, the cells were dehydrated sequentially in 20%, 50%, 70%, 100% ethanol (anhydrous) for 5 min each, followed by anhydrous ethanol at RT for twice, each for10 min. The cells were infiltrated with Epon-Aradite (Ted Pella, Redding, CA) (50% for 1h and 100% for 1 day with 2 changes). Samples were polymerized at 60 °C for two days. Ultrathin sections (about 90 nm) were cut on Leica EM UC6 Ultramicrotome (Leica, Buffalo Grove, IL) (Leica, Buffalo Grove, IL) and collected on copper slot grids. The sections were examined under a FEI Tecnai12 transmission electron microscope (FEI, Hillsboro, Oregon) operating at a beam energy of 120keV. Images were acquired by using a Gatan 2k x 2k cooled CCD camera (Gatan, Warrendale, PA).

#### Image analysis

TEM images at 6500X magnification were used to create binary masks of mitochondria using empanada-napari ^77^. These binary masks were further analyzed using Python libraries. Binary masks were loaded using the `tifffile` library and visualized using `matplotlib.pyplot` to ensure correct loading and preprocessing. The binary images were labeled and region properties extracted using the `skimage.measure` library. Only regions with an area above a defined threshold (10,000 pixels) were considered for analysis. The extracted properties included area, perimeter, convex area, and solidity for each valid mitochondrion. Statistical significance was assessed with a Mann-Whitney U test. Differences in properties between the samples was conducted at the per-mito and per-cell level.

### Puromycin-based FACs analysis of protein translation

Primary samples were collected under IRB approved protocols with written informed consent. Bone marrow mononuclear cells were isolated using the Ficoll-Paque method. Briefly, bone marrow aspirates were diluted 1:2 with PBS and overlayed on Ficoll at a 1:1 volume ratio then spun at 3000 rpm for 15 min. The mononuclear layer was removed, and RBCs were lysed with Qiagen RBC lysis solution. Cells were then washed and counted for downstream use. Protein translation analyses were adapted from the previously published SCENITH and RiboPuromycylation assays ^46,78,79^. Human bone marrow cells isolated from primary samples were resuspended in RPMI 1640 media supplemented with 10% fetal bovine serum (FBS), Glutamax, and PenStrep (R10 Media).

For drug treatments, puromycin stock solution (10 mg/mL) was diluted to a final concentration of 10 µg/mL (Sigma P4512-1MLX10), 2-Deoxy-D-glucose (2DG) stock solution (2 M) was diluted to a final concentration of 150 mM (Sigma D8375-5G), oligomycin stock solution (10 mM) was diluted to a final concentration of 1 µM (Sigma 04876), and cycloheximide stock solution (10 mg/mL) was diluted to a final concentration of 100 µg/mL (Sigma 239764-1GM). R10 media containing these drugs and puromycin (R10P-Drugs) were prepared at 2x concentrations.

Bone marrow cells were resuspended at a concentration of 4 x 10^6 cells/mL in warm R10 media and plated in 50 µL volumes (0.2 x 10^6 cells/well) in 96-well plates for each condition: DMSO, 2DG, Oligomycin, 2DG + Oligomycin (DGO), Cycloheximide (CHX), and no Puromycin. Cells were incubated at 37°C for 1 hour. Subsequently, 50 µL of R10-Drug solution (2x) was added to each well and incubated for 15 minutes at 37°C, followed by the addition of 100 µL of R10P-Drug solution (2x) and a further incubation at 37°C for 30 minutes. Cells were then centrifuged at 4°C, washed with 250 µL of cold PBS, and centrifuged again at 4°C.

For staining, cells were incubated with 50 µL of Live Dead Violet (1:1000 dilution) for 15 minutes in PBS. Cells were washed with PBS containing 2% FBS and then stained with surface antibodies, followed by another wash with PBS containing 2% FBS. Cells were fixed and permeabilized using the eBioscience FoxP3 Staining Buffer Set (Thermo Fisher, Ref 00-5523-00) according to the manufacturer’s instructions. Briefly, cells were incubated with 100 µL of Fix/Perm solution at 4°C for at least 60 minutes, washed with 100 µL of Perm Buffer, and then resuspended in 50 µL of Perm Buffer. Subsequently, cells were stained with 0.5 µL of AlexaFluor647-conjugated anti-puromycin antibody; the anti-PMY-2A4 Ab was directly conjugated (using Life Technologies protein labeling kit as per the manufacturer’s instructions) and the. () in 50 µL of Perm Buffer and incubated at 4°C for 60 minutes. After staining, cells were washed with PBS containing 2% FBS. Finally, the stained cells were analyzed using flow cytometry [BD LSRFortessaTM Cell Analyzer; # 649225] to quantify puromycin incorporation, which serves as a proxy for cellular translation activity and energy production. This assay allowed us to assess the impact of various metabolic inhibitors on translation in primary bone marrow samples. Mitochondrial dependence was calculated as in ^45^.

| Antibody/Fluorochrome | Company Ref |
| --- | --- |
| **CD34 FITC** | BD 555821 |
| **CD11b PE** | Beckman Coulter IM2581U |
| **GlycophorinA BV421** | BD 562938 |
| **Anti-Puromycin AlexaFluor647 PMY-2A4** | RRID:AB_2619605 |
| **Anti-Puromycin 2A4 AlexaFluor647** | Biolegend 381507 |
| **LIVE/DEAD™ Fixable Near-IR: Dead Cell Stain Kit** | Thermo Fisher L10119 |

### CD34+ cell isolation

CD34+ cells were isolated from ficolled bone marrow mononuclear cells using the EasySep™ Human CD34 Positive Selection Kit II (STEMCELL Technologies, 17856) according to the manufacturer’s instructions in PBS containing 2% fetal bovine serum and 1 mM EDTA. To improve recovery, the addition of RapidSphere particles was increased to 100 µL/mL in step 4, and the magnetic separation process was reduced to four 3-minute washes. CD34+ cells were then suspended in the indicated media and prepared for use.

### 2-color smFISH (CD34 + cells)

A Cellvis P96-1.5H-N plate was coated with a 1:10 dilution of poly-L-lysine in sterile water (Thermo Fisher, P8920). Approximately 50-100 µl of the dilution was added to each well, left for 10 minutes, and then the solution was removed carefully to avoid scraping the applied coat. The plate was air-dried with the lid on in a sterile hood for a minimum of 2 hours. CD34+ cells were plated at a density of approximately 25,000-100,000 cells per well and spun down at 250 RPM for 3 minutes in StemSpan SFEM II media (STEMCELL Technologies, 09655). Pre-warmed media was used to resuspend the CD34+ cells, which were then filtered to remove debris before plating. After plating, cells were incubated for 1 hour at 37 °C to recover from Ficoll and CD34+ isolation and to attach to the plate. The cells were fixed with 100 µl of 4% paraformaldehyde in DPBS for 15 minutes at room temperature, followed by three washes with 1X DPBS for 10 minutes each at room temperature. The fixed cells were stored in 70% ethanol at 4°C for up to 1 month with the plate sealed using adhesive foil cover. Optimal results were observed when plates were stored in 70% ethanol at 4°C between 2-7 days before staining.

For smFISH, cells were washed once with 1X DPBS for 10 minutes at room temperature, then incubated with smFISH wash buffer (20% Formamide, 2X SSC) for 45 minutes at room temperature. The wash buffer was removed completely, and hybridization buffer (20% Formamide, 2X SSC, 10% dextran sulfate) with probes (100-125 nM) was added to the cells. The total volume of hybridization buffer with probes was 50 µl. The plate was incubated at 37°C for 4 hours, sealed with foil cover. After hybridization, cells were washed twice with pre-warmed RNA-FISH wash buffer for 35 minutes at 37°C, followed by a wash with 2X SSC for 10 minutes at room temperature, and another wash with 1X DPBS for 5 minutes at room temperature. Cells were then stained with DAPI (0.5 µg/mL) for 2 minutes at room temperature, followed by two washes with 1X DPBS for 10 minutes each at room temperature. Finally, 100 µl of 1X DPBS was added per well, and the plate was sealed with adhesive cover. Plates are ready to image and could be stored at 4°C for up to a week, although image quality started to decline after 2 days.

#### smFISH Imaging

Confocal images of CD34+ smFISH were collected on an Airyscan Zeiss LSM880 confocal microscope using the Airyscan module to enhance resolution in XYZ (120 nm x 120 nm x 380 nm) beyond the diffraction limit at 16-bit image depth. Z-stack images were acquired using an Alpha P-Apo 100X/1.46 NA objective with a z-step of 500 nm, consisting of 12-14 z-planes in the stack. The pixel size was set to 71 nm, and the pixel dwell time was 14 µs. Images were acquired sequentially in the following channels: Red (Ex 561 nm, 50% laser power; Em BP 570-620 nm), Far-Red (Ex 633 nm, 50% laser power; Em LP 645 nm), and Blue (Ex 405 nm, 2% laser power; Em BP 420-500 nm). Data collection was controlled using ZEN software on a Windows 10-based system.

#### smFISH Analysis

CD34+ cells were analyzed using the same spatial point pattern analysis methods as described for HBECs (see *2-color smFISH (HBEC cells)*). The main difference for the CD34+ cell analysis was the use of the maximum deviation of the L function from the expected value under CSR as the summary statistic to quantify the degree of clustering or dispersion. Additionally, due to the lower number of CD34+ cells analyzed, we performed 1000 simulations to generate the confidence envelope, compared to 250 simulations for the HBEC analysis. The maximum deviation was calculated for each cell and used to compare spatial patterns across cells from different sample groups. We then plotted the cumulative distribution function (CDF) of the maximum deviations and performed a Mann-Whitney U test to assess statistical significance.

