## Supplemental Data 1 for "U2AF regulates the translation and localization of nuclear-encoded mitochondrial mRNAs"

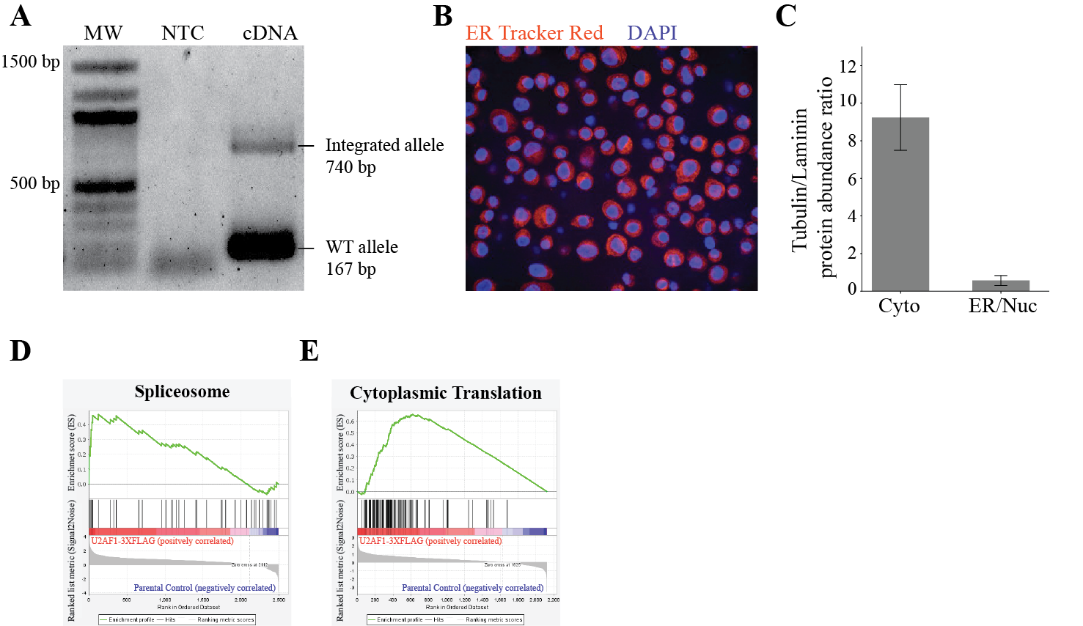

_­_**Fig. S1.** **U2AF1 CRISPR Editing Validation and IP-Mass Spec**

(**A**) 1% agarose gel showing PCR products with lanes labeled at the top. Lane MW: molecular weight ladder (sizes in bp on the left). Lane NTC: no template control, confirming no contamination. Lane cDNA: PCR from U2AF1-FLAG line cDNA used in IP-mass spectrometry. The 740 bp band represents the integrated U2AF1-FLAG allele, and the 167 bp band represents the wild-type allele, as indicated on the right.

(**B**) ­Immunofluorescence image showing isolated nuclei from the ER-nuclear fraction of the IP-mass spectrometry experiment. Nuclei are stained with DAPI (blue) and ER is labeled with ER-Tracker Red (red), confirming the attachment of ER to the nuclei.

(**C**) Tubulin/Laminin protein abundance ratio in cytoplasmic and ER/nuclear fractions from IP mass-spectrometry experiment. The tubulin/laminin ratio was calculated for each replicate from *U2AF1^wt/wt-3XFLAG^* cells, and the mean ratio is presented with error bars indicating the standard deviation. This ratio illustrates the sub-fractionation quality, with a higher tubulin/laminin ratio in the cytoplasmic fraction and a lower ratio in the ER/nuclear fraction.

(**D**) GSEA plot from the cytoplasmic fraction, showing significant enrichment of the 'Splicing' category in *U2AF1^wt/wt-3XFlag^* cells compared to the parental control, as determined by the KEGG gene set.

(**E**) GSEA plot from the ER-nuclear fraction showing significant enrichment of the Cytoplasmic Translation' category in *U2AF1^wt/wt-3XFlag^* cells compared to the parental control, as determined by the KEGG gene set.

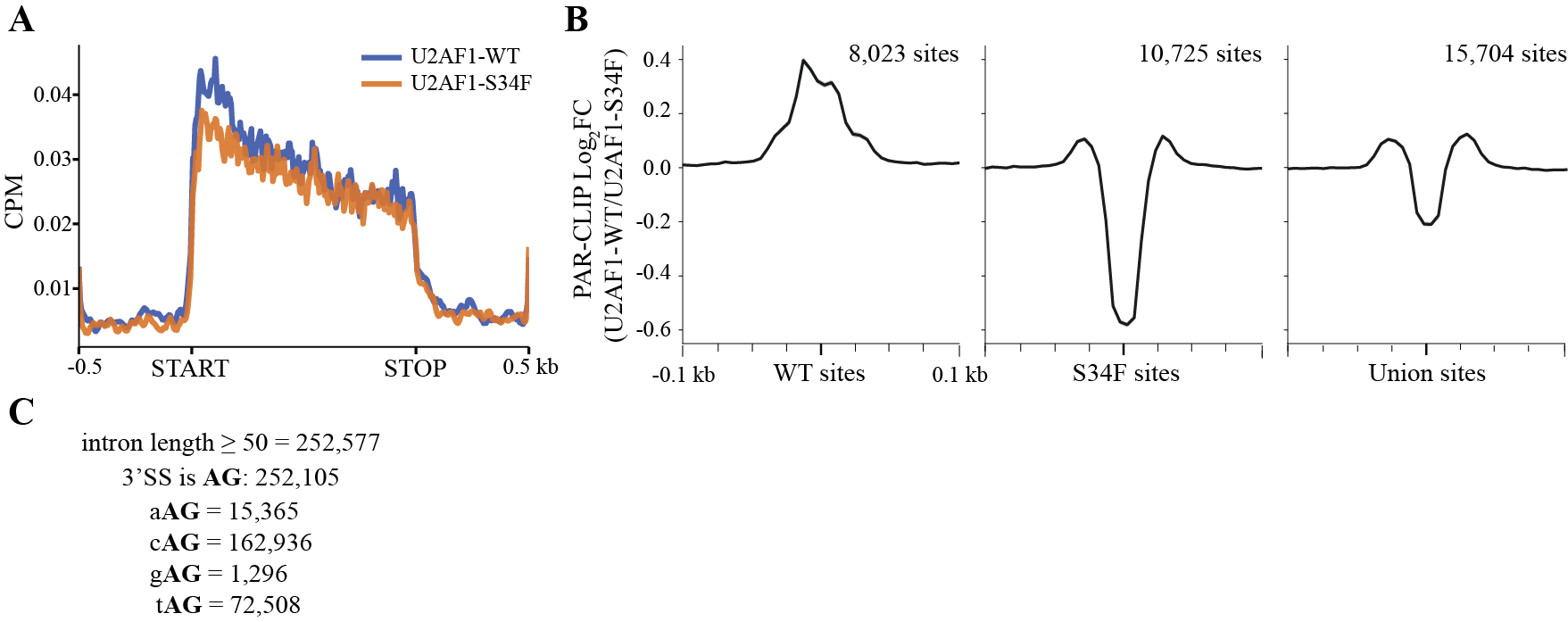

**Fig. S2.** **U2AF Cytoplasmic PAR-CLIP Analysis**

(**A**) Meta-transcript analysis of U2AF1 binding on mRNAs, displaying normalized U2AF1 footprint reads across identified U2AF1 targets for the U2AF1-WT (blue) and U2AF1-S34F (orange) cytoplasmic PAR-CLIP datasets. Binding distribution is shown from 500 bp upstream to 500 bp downstream of the translation start site (START) to the stop codon (STOP). Nucleotide distances in kilobases (kb).

(**B**) Log_2_ fold changes (U2AF1-WT/U2AF1-S34F) in U2AF1 binding across 8,023 WT sites, 10,725 S34F sites, and 15,704 union sites, calculated using cytoplasmic U2AF1 PAR-CLIP datasets. Peaks are centered and extended 100 bases upstream and downstream.

(**C**) Analysis of annotated introns (≥50 bp) from the hg19 genome, totaling 252,577 introns. Among these, 252,105 introns have a 3' splice site containing AG (U2AF1 binding site). The tri-nucleotide sequence nAG was further categorized. Notably, gAG occurs in less than 1% of annotated splice sites.

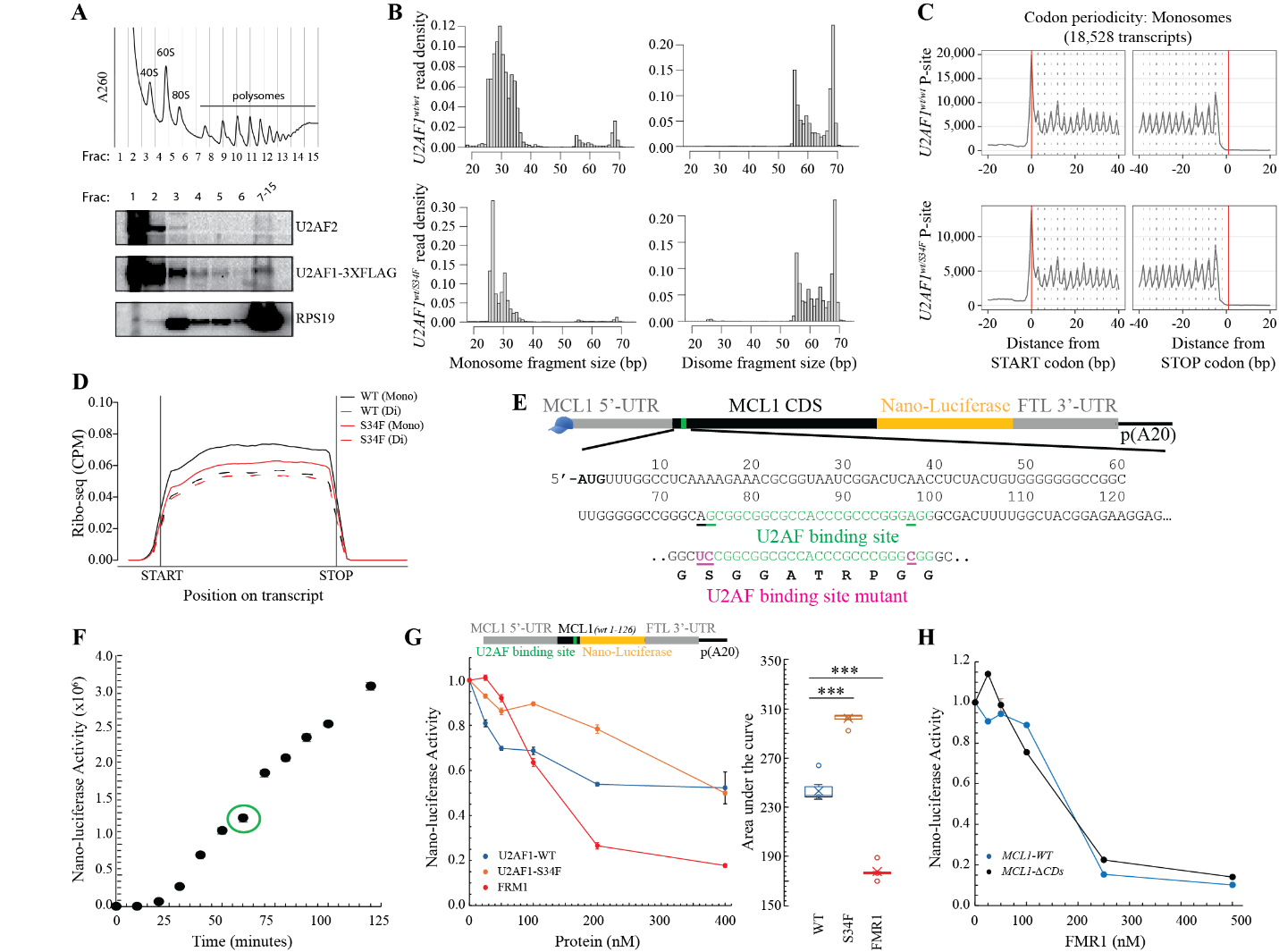

**Fig. S3.** **Multi-experimental analysis of U2AF translational regulation.**

(**A**) Polysome profile followed by western blot analysis from of *HBEC^wt/wt-3XFLAG^* cells. Top: The absorbance at 260 nm (A260) is plotted against the fraction number. The 40S, 60S, 80S (monosome), and polysomes are labeled on the plot. Bottom: Western blot analysis of the polysome profile fractions. Fractions were collected from the polysome profile above and probed for U2AF1-3XFLAG, U2AF2, and RPS19 (a small ribosomal subunit protein).

(**B-D**) Quality control plots for Ribo-sequencing experiments in HBEC *U2AF1^wt/wt^* and *U2AF1^wt/S34F^* cells. (**B**) Fragment size distribution in Ribo-sequencing experiments. Histograms showing the normalized read density (y-axis) versus transcriptome aligned fragment size (x-axis) for monosome and disome ribo-sequencing experiments. The top panels represent *U2AF1^wt/wt^* cells, with monosome fragments on the left and disome fragments on the right. The bottom panels represent *U2AF1^wt/S34F^* cells, with monosome fragments on the left and disome fragments on the right. (**C**) Codon periodicity analysis of monosome Ribo-sequencing. Codon periodicity plots showing the P-site read density around the translation start (left) and stop (right) codons for 18,528 transcripts. The top panels represent *U2AF1^wt/wt^* cells, while the bottom panels represent *U2AF1^wt/S34F^*. The red vertical lines indicate the positions of the start and stop codons. (**D**) Meta-transcript coverage plot from Ribo-sequencing experiments. Meta-transcript coverage plot showing CPM (counts per million) normalized reads for monosomes (solid lines) and disomes (dashed lines) in *U2AF1^wt/wt^* (black) and *U2AF1^wt/S34F^* (red) cells. The x-axis represents the position on the transcript, with the start and stop codons indicated. The plot demonstrates no positional bias in the coverage.

(**E-H**) MCL1 *in vitro* translation assay: timepoint validation and specificity analysis. (**E**) Schematic of the MCL1 nano-luciferase reporter construct (*MCL1-WT*). The schematic illustrates the MCL1 nano-luciferase reporter construct, highlighting the nucleotide sequences starting from the annotated AUG start codon. The U2AF binding site identified by PAR-CLIP is shown in green. The sequences mutated in the *MCL1-mtAGs* reporter construct are indicated in magenta. (**F**) Nano-luciferase activity (y-axis) measured over time (x-axis) using the *MCL1-WT* reporter construct. The 60-minute timepoint, circled in green, indicates the selected timepoint for all in vitro translation assays and falls within the linear range of the assay response. (**G**) *In vitro* translation reporter assay with MCL1 constructs truncated downstream of the U2AF binding site (*MCL1-WT:1-126bp*). Dose-response plot comparing U2AF1-WT, U2AF1-S34F, and FMR1 (non-specific control) RNA binding proteins. The right panel shows the area under the curve (AUC) analysis to highlight significant differences. Error bars represent the standard error of the mean (SEM) and statistical significance was determined using a t-test. **p* < 0.05, ** *p*< 0.01, ****p* < 0.001. (**H**) Dose-response plot shows relative nano-luciferase activity in the presence of increasing concentrations of FMR1 protein. Constructs tested include *MCL1-WT* and *MCL1-ΔCDS* (coding sequence deletion).

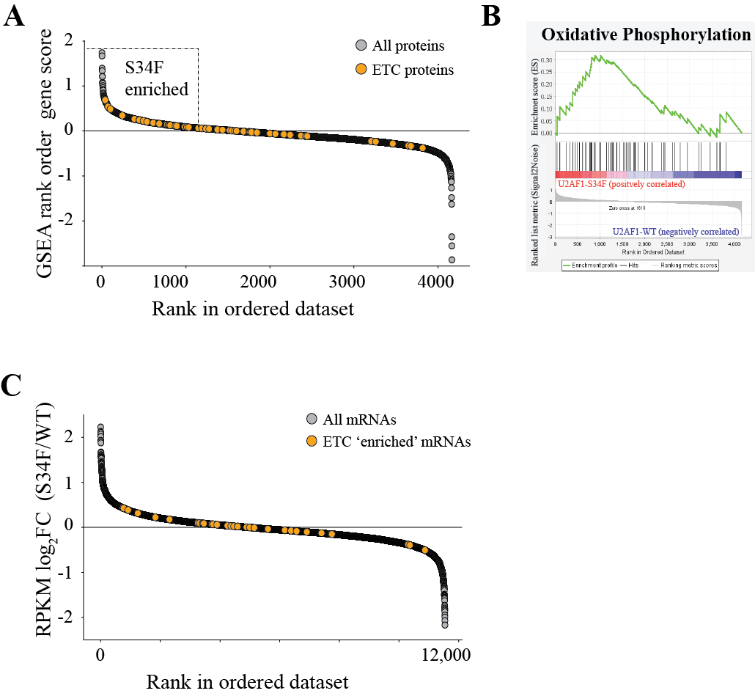

**Fig. S4.** **Global mass spectrometry and RNA-Seq analysis of HBEC *U2AF1^wt/wt^* and *U2AF1^wt/S34F^* cells from previously published datasets**

(**A**) GSEA rank order gene score plot for all proteins, with electron transfer chain (ETC) proteins highlighted in orange. This plot shows that ETC proteins are specifically enriched in the S34F line, as indicated by their higher rank order scores compared to the WT.

(**B**) GSEA of previously published global mass spectrometry data from HBEC *U2AF1^wt/S34F^* versus *U2AF1^wt/wt^* cells, highlighting significant enrichment for the KEGG Oxidative Phosphorylation pathway.

(**C**) Previously published RNA-seq data from HBEC U2AF1-S34F versus WT cells, showing the rank order of log2 fold change (S34F/WT) for all mRNAs. ETC mRNAs that were S34F enriched at the protein level in panel (**A)** are highlighted in orange. This plot demonstrates that the increase in mitochondrial proteins observed in the S34F line is not reflected at the mRNA level.

**Table S3.** Fisher's exact test results for PAR-CLIP datasets of EIF3, G3BP2, and FMR1 in relation to U2AF1 (PAR-CLIP) and annotated mitochondrial mRNAs (MitoCarta 3.0).

|  | **FMR1+** | **FMR1-** | **U2AF1+** | **U2AF1-** | **Mito+** | **Mito-** | **EIF3+** | **EIF3-** |
| --- | --- | --- | --- | --- | --- | --- | --- | --- |
| **FMR1+** |  |  | 3.045 | 0.73 | 0.663 | 1.019 | 2.003 | 0.981 |
| **FMR1-** |  |  | 0.848 | 1.024 | 1.021 | 1.002 | 0.924 | 1.005 |
| **U2AF1+** |  |  |  |  | 1.503 | 0.976 | 3.796 | 0.933 |
| **U2AF1-** |  |  |  |  | 0.993 | 1.006 | 0.631 | 1.01 |
| **Mito+** |  |  |  |  |  |  | 1.595 | 0.99 |
| **Mito-** |  |  |  |  |  |  | 0.972 | 1.002 |
| **EIF3+** |  |  |  |  |  |  |  |  |
| **EIF3-** |  |  |  |  |  |  |  |  |

Table S5. Demographic and disease information of donor and patient samples used in the study.

| Sample | Gender | Age | Diagnosis | NGS | Experimental Assay |
| --- | --- | --- | --- | --- | --- |
| HD2 | F | 75 | Healthy Donor |  | Puromycin-based translation assay |
| HD21 | F | 64 | Healthy Donor |  | Puromycin-based translation assay |
| HD22 | M | 71 | Healthy Donor |  | Puromycin-based translation assay |
| MDS_1 | F | 65 | MDS-IB2 | U2AF1, ASXL1, CREBBP | Puromycin-based translation assay |
| MDS-9 | M | 87 | MDS-LB | DNMT3a, TET2, PTEN | Puromycin-based translation assay |
| MDS-15 | M | 77 | MDS-SF3B1 | DNMT3a, SF3B1 | Puromycin-based translation assay |
| HD23 | F | 64 | Healthy Donor |  | two-color smFISH |
| HD25 | F | 76 | Healthy Donor |  | two-color smFISH |
| HD26 | F | 66 | Healthy Donor |  | two-color smFISH |
| MDS16 | F | 57 | MDS-LB | U2AF1-S34F | two-color smFISH |
| MDS17 | M | 72 | MDS-LB | IDH1, SRSF2 | two-color smFISH |
| AML019 | F | 66 | AML with mutated TP53 | TP53, GATA2, WT1 | two-color smFISH |

_HD = Healthy donor, MDS = Myelodysplastic syndromes, AML = Acute myeloid leukemia, and NGS = Nucleotide_
